# SMCHD1 Promotes ATM-dependent DNA Damage Signaling and Repair of Uncapped Telomeres

**DOI:** 10.1101/669119

**Authors:** Aleksandra Vančevska, Verena Pfeiffer, Marianna Feretzaki, Wareed Ahmed, Joachim Lingner

## Abstract

SMCHD1 (structural maintenance of chromosomes flexible hinge domain containing protein 1) has been implicated in X-chromosome inactivation, imprinting and DNA damage repair. Mutations in *SMCHD1* can also cause facioscapulohumoral muscular dystrophy. More recently, SMCHD1 has also been detected as component of telomeric chromatin. Here, we identify requirements of SMCHD1 for DNA damage signaling and non-homologous end joining (NHEJ) at unprotected telomeres. Co-depletion of SMCHD1 with TRF2 reduced the rate of 3’ overhang removal in time course experiments and the number of telomere end fusions. In SMCHD1 deficient cells, the formation of ATM pS1981, γH2AX and 53BP1 containing telomere dysfunction induced foci (TIFs) were diminished indicating defects in checkpoint signaling. Strikingly, removal of TPP1 and subsequent activation of ATR signaling rescued telomere fusion events in TRF2-depleted *SMCHD1* knockout cells. Together, these data indicate that SMCHD1 depletion reduces telomere fusions in TRF2-depleted cells due to defects in ATM-dependent DNA checkpoint signaling. SMCHD1 mediates DNA damage signaling activation upstream of ATM phosphorylation at uncapped telomeres.

## Introduction

Arguably the most fundamental function of telomeres is to suppress at chromosome ends DNA damage signaling and DNA end repair (Muller, 1938; McClintock, 1941). This is achieved through the recruitment of specialized proteins that bind directly or indirectly to telomeric repeat DNA, which consists of hundreds to thousands of 5’-TTAGGG-3’/5’-CCCTAA-3’ repeats in vertebrates. Most abundant at telomeres are the shelterin proteins comprising TRF1, TRF2, RAP1, TIN2, TPP1 and POT1 (de Lange, 2018; Lazzerini-Denchi & Sfeir, 2016). TRF1 and TRF2 bind as homodimers to the double stranded telomeric DNA repeats. Depletion of TRF2 from chromosome ends occurs naturally upon telomere shortening in senescent cells (Karlseder *et al*, 2002; Cesare *et al*, 2013). TRF2 depletion leads to ATM kinase activation and a long-lasting DNA damage response (DDR) promoting cellular senescence (Denchi & de Lange, 2007) or apoptosis (Karlseder *et al*, 1999). Inactivation of the DDR in senescent cells occurs during tumorigenesis (Shay & Wright, 2011; Maciejowski & de Lange, 2017). The ensuing cell proliferation leads to further telomere shortening and further TRF2 depletion culminating in telomere crisis in which chromosome ends are fused to one another by alternative nonhomologous DNA end joining (alt-NHEJ), which relies on DNA ligase 3 and poly(ADP-ribose) polymerase 1 (PARP1) (Jones *et al*, 2014). Experimental depletion of TRF2 in cells with normal telomere length also leads to ATM-dependent DDR activation and telomere end joining, which in this case is mediated by the classical NHEJ pathway involving DNA ligase 4 and the KU70/80 heterodimer (Celli & de Lange, 2005). Significantly, classical NHEJ at TRF2-depleted telomeres requires DDR activation (Denchi & de Lange, 2007).

The DDR promotes genome stability regulating DNA repair, chromatin remodeling, transcription, cell cycle arrest, senescence and apoptosis (Ciccia & Elledge, 2010; Panier & Durocher, 2013). DDR activation at DNA double strand breaks and uncapped telomeres involves ATM recruitment to chromatin by the MRE11/RAD50/NBS1 (MRN) complex, which also promotes conformational changes stimulating ATM kinase activity (Paull, 2015). In addition to the interaction with NBS1 in the MRN complex, ATM activation also depends on Tip60/KAT5-dependent acetylation of K3016 in ATM (Sun *et al*, 2009). Active ATM leads to autophosphorylation at S1981 and phosphorylation and activation of hundreds of downstream DDR substrates (Matsuoka *et al*, 2007), such as the CHK2 kinase, p53, NBS1, 53BP1 and H2AX.

SMCHD1 is a non-canonical member of the structural maintenance of chromosomes (SMC) protein family (Blewitt *et al*, 2008), which includes among others the SMC1/3 cohesion and SMC2/4 condensin complex components, the SMC5/6 complex which is involved in homologous recombination and RAD50. As other SMC proteins, SMCHD1 contains a hinge domain flanked by coiled-coil domains. However, unlike SMC1-6, SMCHD1 forms homodimers (Brideau *et al*, 2015). Furthermore, it contains a GHKL (gyrase, Hsp90, histidine kinase, MutL)-type ATPase rather than the bipartite ABC-type ATPase domain typically seen in SMC proteins (Brideau *et al*, 2015). Several studies implicate roles of SMCHD1 in modulating chromosome architecture at the inactive X chromosome and at Hox clusters (Nozawa *et al*, 2013; Wang *et al*, 2018; Jansz *et al*, 2018; Gdula *et al*, 2019). SMCHD1 mediates the compaction of the inactive X chromosome in females linking the H3K9me3 and the XIST-H3K27me3 domains (Nozawa *et al*, 2013). Through mediating chromatin interactions on the inactive X chromosome, SMCHD1 may also promote chromatin mixing and drive attenuation of chromosomal compartments and topologically associated domains (TADs) (Wang *et al*, 2018). SMCHD1 localization to the inactive X chromosome in mouse cells is dependent on a pathway involving the *Xist* long noncoding RNA, Hnrnp K and PRC1 (Jansz *et al*, 2018). Alternative mechanisms of SMCHD1 recruitment to chromatin may involve H3K9me3, HP1γ and affinity for nucleic acids (Nozawa *et al*, 2013). Apart from binding the inactive X chromosome, SMCHD1 is also recruited to sites of DNA damage induced by laser micro-irradiation (Coker & Brockdorff, 2014) or zeocin drug treatment and it has been implicated in promoting DNA repair by NHEJ over homologous recombination (Tang *et al*, 2014). Finally, SMCHD1 has been detected in proteomic analyses of telomeric chromatin (Déjardin & Kingston, 2009; Grolimund *et al*, 2013; Bartocci *et al*, 2014). Specifically, SMCHD1 was enriched at telomeres that were overly long and showed a lower density of TRF2 (Grolimund *et al*, 2013). However, the roles of SMCHD1 at telomeres remained enigmatic.

Here, we discover critical functions of SMCHD1 at telomeres that were deprived of TRF2. Significantly, depletion of SMCHD1 impairs with ATM-dependent DNA damage signaling at TRF2-depleted telomeres. At the same time telomere end fusions were diminished indicating crucial roles of SMCHD1 in DNA damage signaling or repair. Experimental activation of the ATR checkpoint at TRF2-depleted telomeres re-instigated chromosome end fusions in the absence of SMCHD1 unraveling a requirement of SMCHD1 for checkpoint activation but not directly the DNA repair reaction. Our data indicate that SMCHD1 is required promoting ATM dependent DDR activation.

## Results

### SMCHD1 is required for efficient telomere-end-to-end fusions at TRF2-depleted telomeres

In previous work we observed in HeLa cells enrichment of SMCHD1 at long telomeres with an average length of 30 kb over telomeres with an average length of 10 kb (Grolimund *et al*, 2013). In addition, over-elongated telomeres showed a lower density of TRF2. We therefore tested if shRNA-mediated TRF2 depletion in HeLa cells is sufficient to enhance association of SMCHD1 with telomeres of normal length. SMCHD1 association with telomeric DNA was assessed by chromatin immunoprecipitation upon which co-precipitated telomeric DNA was detected by Southern hybridization. Indeed, immunoprecipitated SMCHD1 was associated with more telomeric DNA in TRF2-depleted cells (Fig EV1). A probe for Alu-repeat DNA served as a negative control.

TRF2-depleted telomeres trigger an ATM-dependent DNA damage response and they undergo NHEJ-mediated telomere end-to-end fusions (Denchi & de Lange, 2007). In order to assess potential roles of SMCHD1 for these processes, we used CRISPR/Cas9 technology to disrupt the *SMCHD1* gene and we developed shRNA vectors for the depletion of SMCHD1 (Fig 1). Three different guide RNAs were used for generating *SMCHD1* knockout clones in HeLa cells and in a HeLa cell clone in which TRF2 could be depleted using an inducible shRNA (Grolimund *et al*. 2013). Individual clones were screened for loss of SMCHD1 protein expression on Western blots using antibodies recognizing SMCHD1 peptides near the N- and C-termini (Fig 1A and Fig EV2A). This analysis suggested complete loss of SMCHD1 protein expression in all three clones (Fig EV2A). Analysis of the knockout clones by PCR amplification of the targeted loci and DNA sequencing revealed introduction of frameshift mutations near the N-terminus of *SMCHD1* leading to premature stop codons (Fig EV2B), which can explain the loss of SMCHD1 protein expression. In addition, two shRNAs mediated efficient depletion of SMCHD1 protein (Fig EV3 and further below). ShRNA-mediated TRF2 depletion during 5 days (Fig 1A) triggered end-to-end fusions at 20% of the chromosome ends as assessed by the analysis of metaphase chromosome spreads (Fig 1B,C). Strikingly, the telomere fusions were reduced to roughly 3-4% when TRF2 was depleted in two different *SMCHD1* knockout clones. Similar results were obtained upon shRNA-mediated co-depletion of SMCHD1 with TRF2 (Fig EV3) confirming critical roles of SMCHD1 for efficient telomere end-to-end fusions upon TRF2 loss. The effects of SMCHD1-loss on telomere end-to-end fusions were not due defects in cell cycle progression as the cell cycle profile was not strongly affected in *SMCHD1* knockout cells versus WT (Fig 1D).

**Figure 1.**
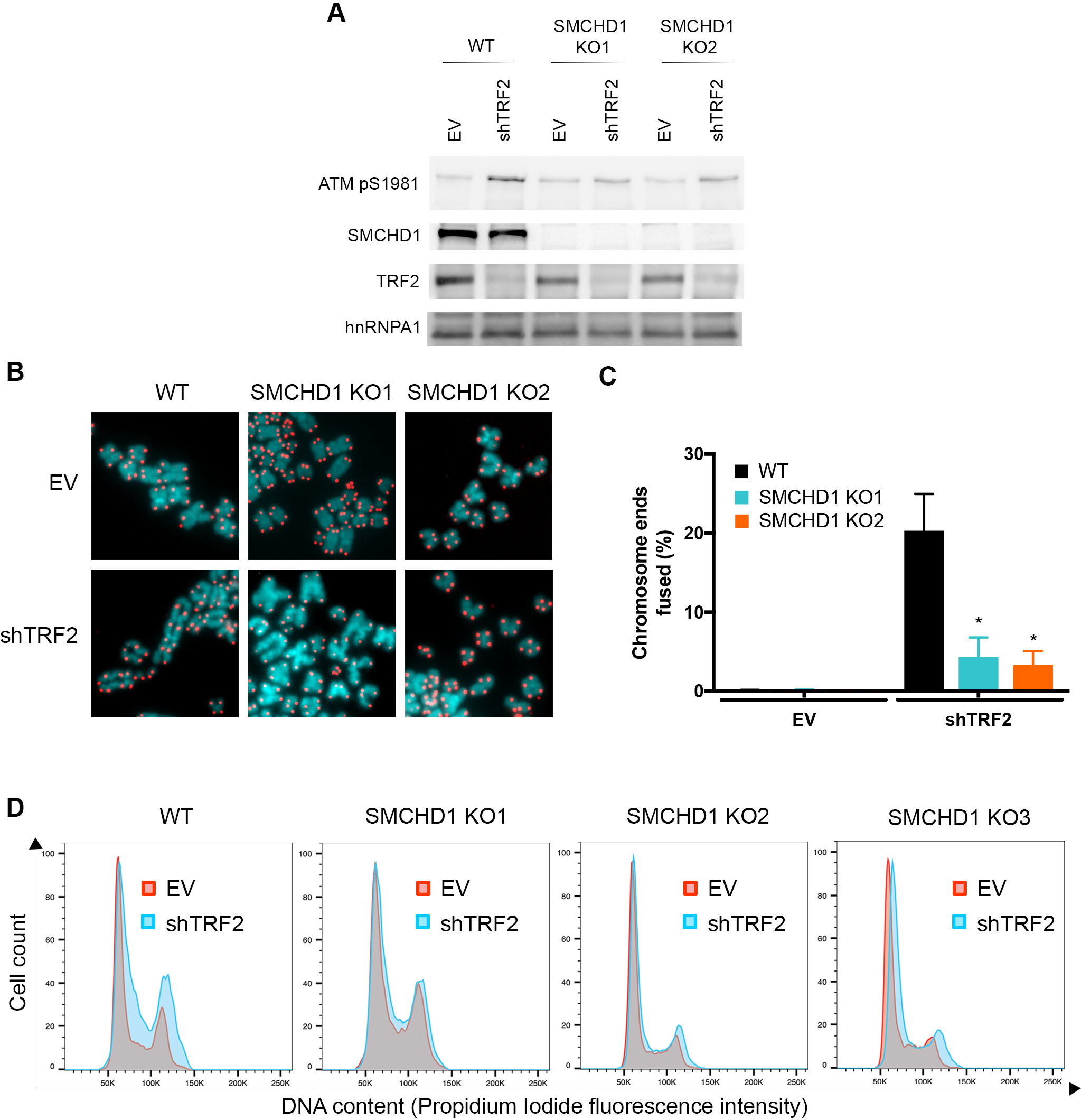
SMCHD1 promotes c-NHEJ at dysfunctional telomeres. (*A*) Western Blot detection of ATM pS1981, SMCHD1, TRF2 and hnRNPA1 loading control in wild-type and two *SMCHD1* knockout (KO1 and KO2) HeLa cells transfected with shTRF2 plasmid or EV control. (*B*) Representative metaphase spreads from wild-type and *SMCHD1* knockout HeLa cells transfected with shTRF2 plasmid or EV control. Telomeric signals were detected with Cy3-OO-(CCCTAA)3 and are false colored in red, DNA is stained with DAPI and is false colored in cyan. (*C*) Quantification of telomere fusions from experiment shown in *D)*. Bars represent average number of fused chromosome ends. SDs were obtained from 3 independent experiments (>3,000 telomeres counted/condition/experiment). (*) P < 0.05; unpaired two-tailed Student’s t-test. (*D*) Representative cell cycle profiles from two biological replicates of WT and SMCHD1 KO HeLa cells transfected with shTRF2 plasmid or EV control. Cells were stained with propidium iodide and analyzed by Flow cytometry.

### SMCHD1 promotes DNA end processing for NHEJ at TRF2-depleted telomeres

Upon TRF2-depletion, telomeric DNA is first processed to remove the 3’ overhang. The blunt end telomeres are then fused by NHEJ (Celli & de Lange, 2005). To better understand the roles of SMCHD1 in these processes, we followed telomeric DNA processing and fusions in time-course experiments in which TRF2 was depleted using an inducible shRNA in *SMCHD1* wild type and knockout cells (Fig 2A). Quantification of telomere end-to-end fusions showed again strong reduction but not abolishment of fusion events in *SMCHD1* knockout cells (Fig 2B). Removal of the telomeric 3’ overhang was assessed by native in gel hybridization in which the radiolabeled probe detects only the telomeric 3’ overhang but not the double stranded telomeric DNA, which remains base-paired (Fig 1C, left panel). As expected the overhang signal was lost upon *in vitro* treatment of the DNA prior to gel loading with Exonuclease 1 from *E. coli* which removes the 3’ overhang (left panel, lanes designated with +Exo). Upon denaturation of the same gel, however, single and double stranded telomeric DNA is detected with the probe (right panel). Inspection of the native gels (Fig 2C, left panel, see short run) and quantification revealed that the *SMCHD1* knockout cells lost the telomeric 3’ overhang considerably slower than the *SMCHD1* wild-type cells (Fig 2C). Furthermore, the signal for fused telomeres which is fully double stranded and therefore can only be detected in the denatured gel (Fig 2C, right panel, see long run), was strongly reduced in the *SMCHD1* knockout cells (compare signal of fused to non-fused telomeres in each lane). These results are consistent with the metaphase chromosome analysis of Figures 1B,C and 2B. During the time course, we also observed a shift of the telomeric signals over time towards longer telomeres which was expected as TRF2 negatively regulates telomere elongation by telomerase (Smogorzewska *et al*, 2000). Telomere elongation was also apparent on the native gel indicating that these telomeres were not fused. Altogether, this analysis indicated that the first step of the telomeric DNA end-fusion reaction, the DNA end processing step was strongly delayed in the absence of SMCHD1.

**Figure 2.**
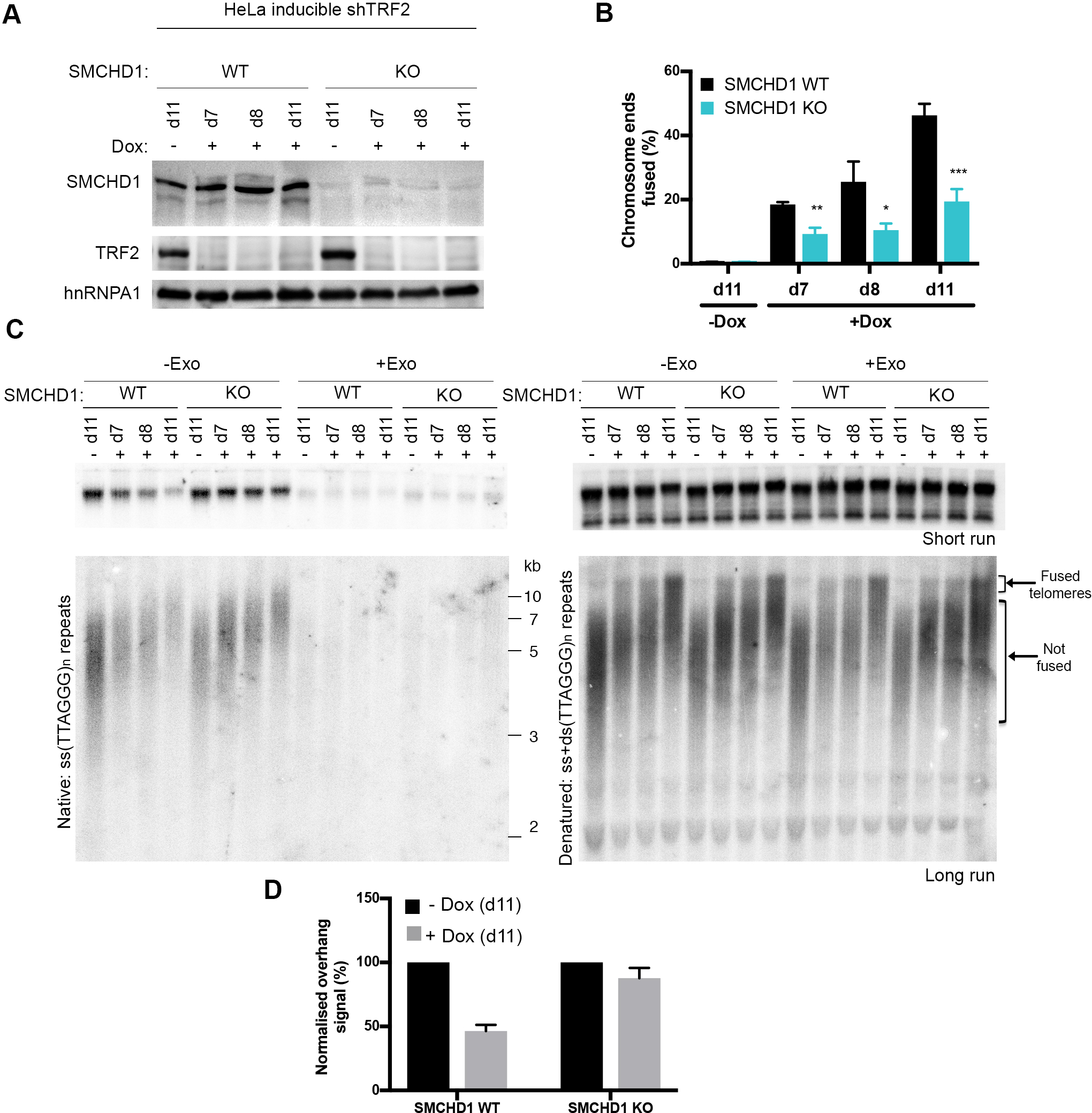
*SMCHD1* loss slows down overhang processing at TRF2 depleted telomeres. (*A*) Western Blot detection of SMCHD1, TRF2 and hnRNPA1 in *SMCHD1* wild-type or *SMCHD1* knockout HeLa inducible shTRF2 cells treated with or without doxycycline for the indicated number of days (d7, d8, d11). (*B*) Quantification of telomere fusions in *SMCHD1* wild-type or *SMCHD1* knockout HeLa inducible shTRF2 cells treated with or without doxycycline for the indicated number of days (d7, d8, d11). Bars represent average number of fused chromosome ends. SDs were obtained from 3 independent experiments (>1,900 telomeres counted/condition/experiment). (***) P < 0.001, (**) P< 0.01; (*) P< 0.01; unpaired two-tailed Student’s t-test. (*C*) Terminal Restriction Fragment (TRF) analysis of telomeric DNA to detect 3’overhang processing of genomic DNA isolated from *SMCHD1* wild-type or *SMCHD1* knockout HeLa inducible shTRF2 cells treated as in the experiment in (*B*). (*Left*) Radiolabeled (CCCTAA)n probe was hybridized with a short run (upper panel) and long run (lower panel) native DNA gel to detect the signal of the telomeric 3’ overhang. Samples used for the short and the long run were from the same digestion split into two. Exo I treatment (+ Exo) was used as a control that single stranded telomeric signal was terminal. (*Right*) The total TTAGGG signal in the same lane was deteced upon denaturation and hybridization with the same probe. (*D*) Quantification of the telomeric overhang signal at d11 after doxycyclin addition to *SMCHD1* wild-type and *SMCHD1* knockout HeLa shTRF2 inducible cells. The bar graph represents the average overhang signal intensity from two biological replicates as percentage of the signal in the cells untreated with doxycycline.

### SMCHD1 promotes ATM activation and DDR at TRF2-depleted telomeres

NHEJ of TRF2-depleted telomeres is strictly dependent on activation of the DDR at uncapped telomeres (Denchi & de Lange, 2007). Therefore, we tested if SMCHD1 is required for checkpoint signaling. As expected, TRF2-depletion during 5 days led to induction of telomere dysfunction induced foci (TIFs) (Takai *et al*, 2003) in which at S1981 phosphorylated ATM (ATM pS1981), phosphorylated H2AX (γH2AX) and 53BP1 accumulate as foci at telomeres (Fig 3). Strikingly, depletion of TRF2 in the two *SMCHD1* knockout clones showed a strong reduction but not abolishment of all TIF markers indicating reduced DDR at TRF2-depleted telomeres in the absence of SMCHD1. Similarly, we observed reduced TIFs in TRF2-depleted cells that had been treated with SMCHD1 shRNAs (Fig EV4). Finally, we observed in Western blots, that ATM pS1981 was reduced in TRF2-depleted *SMCHD1* knockout cells (Fig. 1A) or upon shRNA-mediated depletion of SMCHD1 (Fig EV3A). Altogether, these results indicate that SMCHD1 is required for efficient ATM activation and the subsequent DDR at TRF2-depleted telomeres. Notably, however, SMCHD1 is not absolutely essential for the DDR. Thus SMCHD1 loss has less severe consequences than MRE11 depletion, which completely abolished DDR and NHEJ at TRF2-depleted telomeres (Fig EV3 and Fig EV4), reminiscent of results obtained in mouse embryonic fibroblasts (MEFs) in which *Mre11* was deleted (Deng *et al*, 2009).

**Figure 3.**
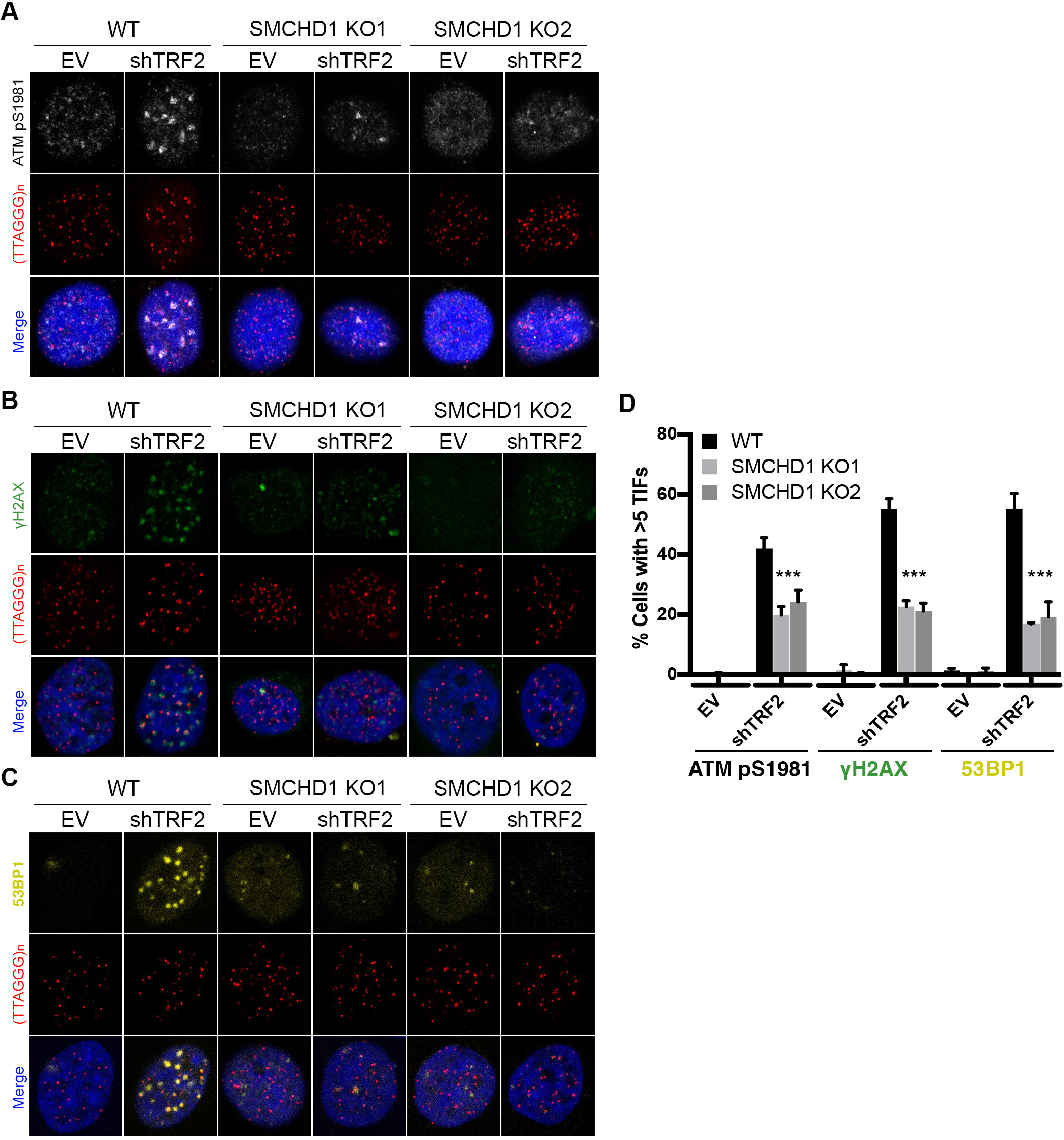
SMCHD1 promotes TIF formation and stimulates ATM signaling from TRF2 depleted telomeres. (*A-C*) Representative images for detection of ATM pS9181, γH2AX, and 53BP1 at telomeres in wild-type (WT) and *SMCHD1* knockout HeLa cells transfected with shTRF2 plasmid and empty vector (EV) control. Immunofluoresence (IF) for ATM pS1981 (gray), γH2AX (green) and 53BP1 (yellow) was combined with telomeric (CCCTAA)3-FISH (red) and DAPI staining total DNA. (*D*) Quantification of the number of cells containing >5 Telomere dysfunction Induced Foci (TIFs) detected as in (*A*)-(*C*). Data represent the mean of 4 independent experiments ± SD (>200 cells/condition/experiment) for ATM pS1981 and 3 independent experiments ± SD (>200 cells/condition/experiment) for γH2AX and 53BP1.

### SMCHD1 is nonepistatic with TERRA and MDC1

The above analyses indicated that SMCHD1 is required for efficient ATM signaling at uncapped telomeres. TRF2 depletion also leads to upregulation of the long noncoding RNA TERRA which is thought to activate the nuclease activity of MRE11 promoting the interaction between MRE11 and its activating lysine demethylase LSD1 (Porro *et al*, 2014b, 2014a). TERRA upregulation occurs independently of ATM and principally, TERRA could act with SMCHD1 upstream of ATM or it could function in a separate pathway. We therefore tested if SMCHD1 is required for the induction of TERRA expression upon TRF2 depletion quantifying TERRA stemming from different chromosome ends by RT-qPCR (Feretzaki & Lingner, 2017) (Fig EV5). The quantification of TERRA demonstrated that TERRA induction upon TRF2-depletion occurs in *SMCHD1*-knockout as well as in WT cells indicating that SMCHD1 is not required for TERRA induction.

Another factor implicated in TIF formation at uncapped telomeres stimulating chromosome end-to-end fusions is MDC1 (Dimitrova & de Lange, 2006). We determined the epistatic relationship of SMCHD1 and MDC1 for TIF formation upon TRF2 loss, depleting MDC1 with shRNAs in WT and *SMCHD1* knockout cells (Fig EV6). As expected TIF formation was reduced upon deletion of *SMCHD1* or depletion of MDC1. Significantly, TIF formation was further reduced upon MDC1 depletion in *SMCHD1* knockout cells unraveling additive effects. Therefore, the two factors behaved in a non-epistatic manner suggesting that SMCHD1 does not directly cooperate with MDC1 for DNA damage signaling. The latter is also consistent with the notion that in contrast to SMCHD1 (Fig 2), MDC1 is not required for efficient removal of the 3’ overhang indicating that it functions further downstream (Dimitrova & de Lange, 2006).

### SMCHD1 is not required for genome-wide ATM signaling

ATM activation at uncapped chromosome ends may involve distinct steps that do not occur upon induction of chromosome breaks elsewhere in the genome. We tested in time course experiments if SMCHD1 is required for ATM activation at chromosome breaks induced by γ-irradiation and if SMCHD1 influences ATM foci disappearance that occurs upon DNA repair (Fig 4). In addition, we followed accumulation and disappearance of DNA damage markers on Western blots. CHK2 which acts downstream of ATM becomes phosphorylated at T68. CHK1 and RPA become phosphorylated by ATR. In the time course experiments, no striking difference was observed between WT and *SMCHD1* knockout cells with regard to foci accumulation and disappearance and the overall accumulation and disappearance of DNA damage markers (Fig 4). Therefore, we conclude that for the bulk of DNA damage sites that were induced by γ-irradiation, SMCHD1 played no crucial role for DNA damage signaling and DNA repair.

**Figure 4.**
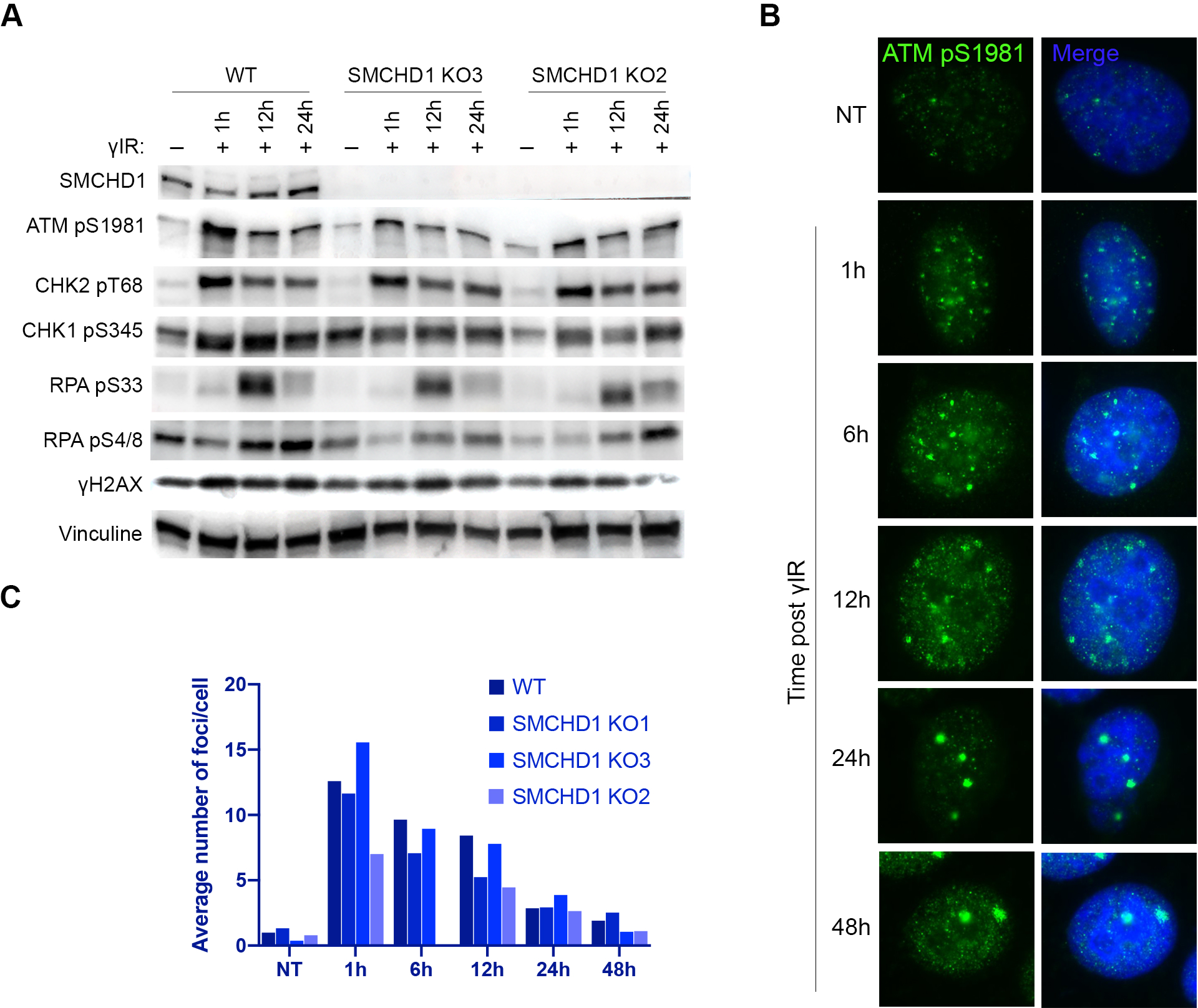
SMCHD1 is not required for genome-wide ATM signaling. (*A*) Western Blot detection of SMCHD1, ATMpS1981, CHK2 pT68, CHK1 pS345, RPA pS33, RPA pS4/8, γH2AX and Vinculin in wild-type or SMCHD1 KO HeLa cells γ-irradiated with 3Gy dose and harvested at the indicated times (1h, 12h, 24h) after the treatment. (*B*) Representative images for Immunofluoresence (IF) detection of ATMpS1981 in wild-type HeLa cells γ-irradiated with 3Gy dose and harvested after the indicated times. (*C*) Quantification of the average number of ATMpS1981 foci per cell detected as in (*B*) in wild-type and SMCHD1 KO γ-irradiated HeLa cells.

### SMCHD1 is required for NHEJ at TRF2-depleted telomeres because of DNA damage response signaling activation

The above results unraveled requirements of SMCHD1 for ATM activation and NHEJ of TRF2-depleted telomeres but they could not distinguish if the effects on NHEJ were solely due to its involvement in checkpoint activation or if SMCHD1 also played direct roles in the DNA processing or end ligation reactions. For addressing this question we were inspired by a previous landmark paper (Denchi & de Lange, 2007), which discovered the requirement of ATM for NHEJ of TRF2-depleted telomeres and which demonstrated that ATM function could be substituted by activated ATR. To activate ATR at telomeres, we depleted TPP1 with shRNAs (Fig 5A), which leads to removal of POT1 from the telomeric 3’ overhang (Frescas & de Lange, 2014). This in turn leads to RPA binding to the single stranded 3’ overhang, subsequent ATR/ATRIP recruitment and checkpoint signaling at chromosome ends (Zou & Elledge, 2003). Significantly, the shRNA-mediated depletion of TPP1 reinstated efficient chromosome end-to-end fusions in *SMCHD1* knockout cells that had been depleted for TRF2 (Fig 5B). Concomitant inhibition of the ATR kinase with an inhibitor (VE-821) (Reaper *et al*, 2011) again prevented efficient end fusions (Fig 5C) indicating that ATR signaling upon TPP1-depletion was responsible for triggering chromosome end-to-end fusions in the absence of SMCHD1.

**Figure 5.**
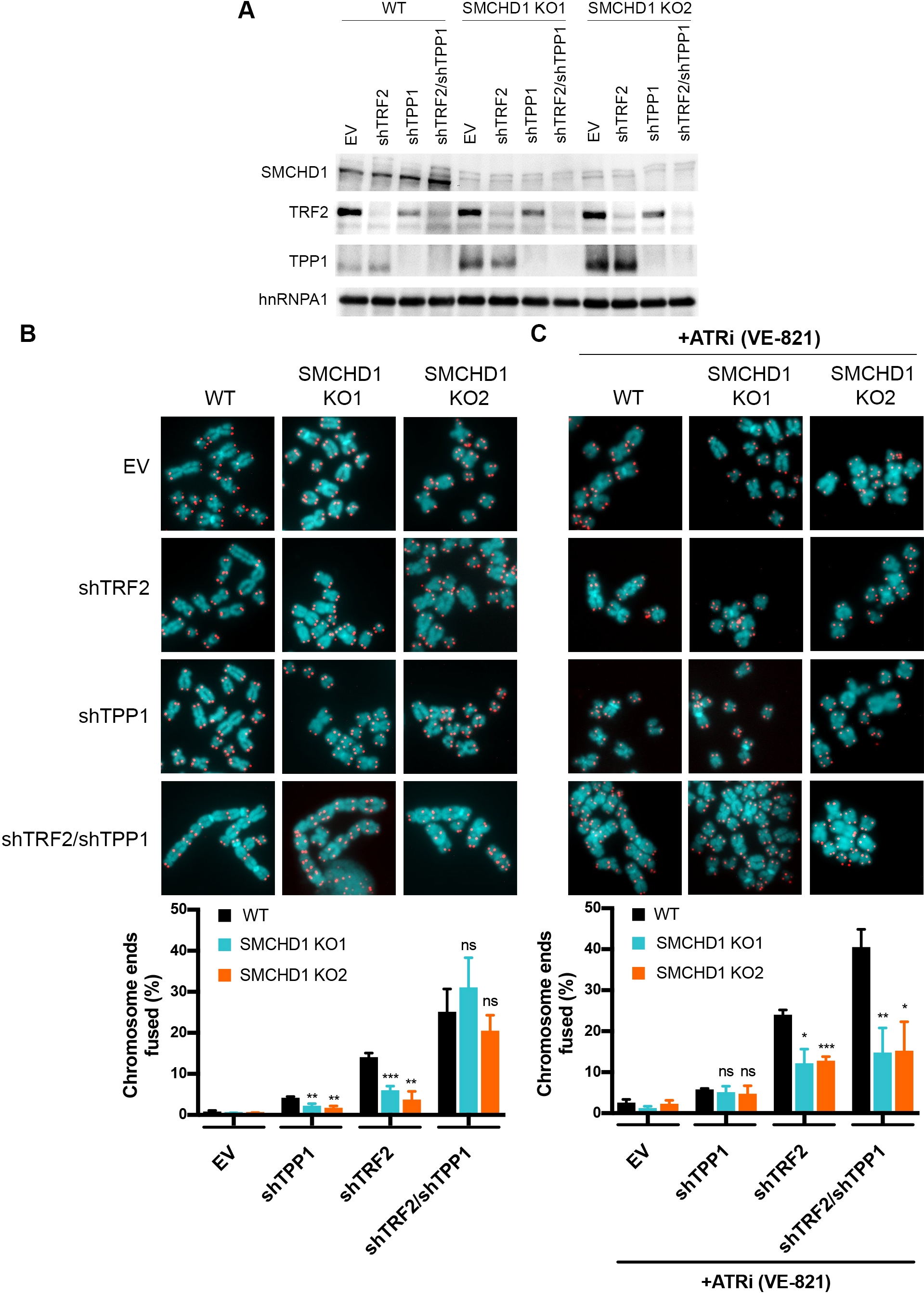
ATR signalling induction by TPP1 removal rescues the telomere fusion defect in *SMCHD1* knockout cells. (*A*) Western Blot detection of SMCHD1, TRF2, TPP1 and hnRNPA1 in wild-type or *SMCHD1* knockout HeLa cells transfected with the indicated shRNAs (shTRF2, shTPP1, shTRF2/shTPP1) or EV control. (*B*) Representative metaphase spreads from HeLa cells transfected with indicated shRNAs or EV control and quantification of telomere fusions. Bars represent average number of fused chromosome ends. SDs were obtained from 3 independent experiments (>2,500 telomeres counted/condition/experiment). (*C*) Representative metaphase spreads from HeLa cells transfected with indicated shRNAs or EV control treated for 4 days with ATRi (VE-821) and quantification of telomere fusions. Bars represent average number of fused chromosome ends. SDs were obtained from 3 independent experiments (>2,000 telomeres counted/condition/experiment).

## Discussion

In this paper we demonstrate that loss of SMCHD1 abolishes efficient DNA damage signaling and NHEJ at telomeres that are depleted of TRF2. The defects of *SMCHD1* knockout cells in signaling and repair can be ascribed to its roles in DDR activation (Fig 6). Indeed, activation of ATR upon depletion of TPP1 was sufficient to suppress the defects of the *SMCHD1* knockout for NHEJ at TRF2-depleted telomeres suggesting that SMCHD1 is required for checkpoint signaling but it is not directly involved in NHEJ. Since SMCHD1 loss in TRF2-depleted cells strongly attenuated ATM activation, our data indicate that SMCHD1 functions in the DDR cascade upstream of ATM phosphorylation (Fig 6). During canonical ATM-dependent DDR at DNA double strand breaks, the MRN complex binds and senses DNA ends recruiting and activating ATM, which then initiates the DNA damage signaling cascade (Paull, 2015). At telomeres, the MRN complex is present even when telomeres are intact (Zhu *et al*, 2000). Indeed, NBS1 of the MRN complex interacts directly with TRF2 but in this context, ATM is not activated (Rai *et al*, 2017). TRF2 inhibits ATM signaling by several mechanisms involving its TRFH and hinge domains (Okamoto *et al*, 2013). The TRFH domain of TRF2 promotes formation of t-loops, which prevents exposure of the chromosome ends to the MRN complex not allowing ATM recruitment or activation (Doksani *et al*, 2013; Van Ly *et al*, 2018). In addition, the TRFH domain of TRF2 interacts at intact telomeres with a non-phosphorylated form of NBS1 preventing ATM activation (Rai *et al*. 2017). Second, through a portion of the hinge domain of TRF2 referred to as iDDR, TRF2 can sever the DDR at the level of the E3 ubiquitin ligase RNF168 which is required for 53BP1 localization to telomeres (Okamoto *et al*, 2013). Upon TRF2 removal, NBS1 is phosphorylated by CDK2 at Ser432 (Rai *et al*, 2017). The t-loops will unwind and MRE11/RAD50 may associate with the uncapped telomeres possibly at their DNA ends in a similar manner as it does with DNA double strand breaks (Syed & Tainer, 2018). Phosphorylated NBS1 may bind to uncapped telomeres via MRE11 enabling ATM recruitment and activation (Rai *et al*, 2017).

**Figure 6.**
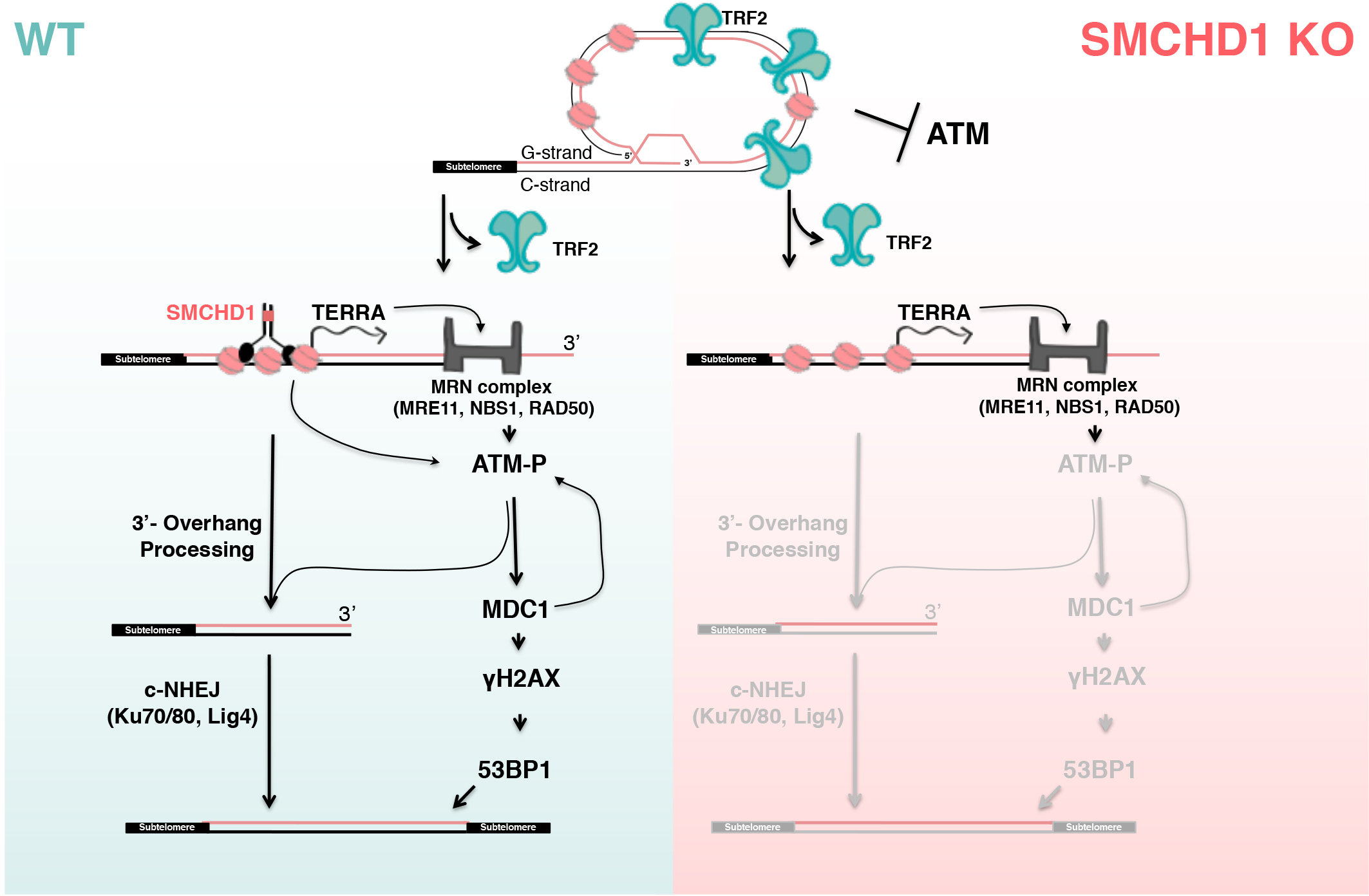
Schematic model of the DNA damage response at uncapped telomeres in *SMCHD1* wild type and knockout cells. Loss of TRF2 leads to t-loop unwinding. In wild type cells, SMCHD1 may remodel the telomeric chromatin which promotes ATM activation by the MRN complex (arrows). ATM activation is required for NHEJ at TRF2-depleted telomeres. In *SMCHD1* knockout cells, ATM activation and DNA damage signaling is attenuated resulting in inefficient 3’ overhang processing and impaired telomere end-to-end fusions. SMCHD1-loss and lack of ATM activation can be compensated for by ATR (not depicted). Only the subset of events and factors are depicted which are directly relevant to this paper.

Our data implicate SMCHD1 at the onset in the damage signaling cascade. We infer that SMCHD1 is not required for t-loop unwinding as its function at uncapped telomeres could be reinstated upon ATR activation (Fig 5) and checkpoint kinases have not been implicated in t-loop unwinding (Van Ly *et al*, 2018). SMCHD1 promotes ATM activation as in its absence ATM is not efficiently phosphorylated and ATM pS1981 does not accumulate substantially in TIFs (Fig 6). SMCHD1 contains an N-terminal ATPase domain and a C-terminal hinge domain mediating homodimerization (Brideau *et al*, 2015). We speculate that SMCHD1 may promote ATP-dependent chromatin remodeling at telomeres upon t-loop unfolding. A role in chromatin architecture and remodeling would be analogous to the functioning of other SMC proteins (van Ruiten & Rowland, 2018) and consistent with the described roles of SMCHD1 in modulating chromosome structure (Nozawa *et al*, 2013; Wang *et al*, 2018; Jansz *et al*, 2018; Gdula *et al*, 2019). For example, SMCHD1 may modulate the telomere structure at TRF2-depleted telomeres at the molecular level to expose telomeric DNA ends and favor MRE11 remodeling thus assisting ATM recruitment and activation (Fig 6). This function of SMCHD1 appears not to involve TERRA induction which occurs upon TRF2-depletion independently of SMCHD1 and ATM. Our epistatic analyses with MDC1 also suggest that SMCHD1 and MDC1 act at separate steps in DNA damage signaling.

At the inactive X chromosome in females, SMCHD1 has been implicated in chromosome compaction linking H3K9me3 rich with H3K27me3 rich domains (Nozawa *et al*, 2013). At telomeres, however, we did not detect notable effects of SMCHD1 depletion on telomere compaction (data not shown). Thus, although H3K9me3 may be important for SMCHD1 binding to uncapped telomeres, SMCHD1-loss does not alter chromatin compaction at telomeres at detectable levels as seen at the inactive X chromosome.

ATM activation is not only required for NHEJ of uncapped telomeres but also for NHEJ of a subset of DNA breaks which occur in heterochromatic regions of the genome (Goodarzi *et al*, 2008). It has been proposed that ATM signaling at DNA breaks temporarily perturbs heterochromatin to promote processing of otherwise inflexible chromatin (Goodarzi *et al*, 2008). We tested if SMCHD1 promotes also ATM activation at DNA breaks induced by γ-irradiation elsewhere in the genome. Our data indicate that SMCHD1 plays no essential role for damage signaling by ATM and DNA repair for the majority of damage sites that were induced by γ-irradiation. Nevertheless, we do not exclude that SMCHD1 participates in damage signaling at other specialized regions of the genome which possibly adopt a heterochromatic structure similar to telomeres. Consistent with this notion are previous observations, which demonstrated recruitment of SMCHD1 to laser-beam induced sites of DNA damage (Coker & Brockdorff, 2014; Tang *et al*, 2014). However, our data do not support a general role of SMCHD1 in ATM activation and DNA repair. They unravel for the first time specific roles of SMCHD1 at uncapped telomeres to trigger damage signaling activation and we determine that its requirement for NHEJ at uncapped telomeres is solely explained by its contribution to the DDR.

ATM activation upon telomere shortening and TRF2 depletion contributes to the induction of cell cycle arrest and cellular senescence (d’Adda di Fagagna *et al*, 2003). Our results implicate SMCHD1 in damage signaling from unprotected telomeres. Mutations in SMCHD1 have been linked to several diseases including facioscapulohumoral muscular dystrophy (FSHD) and Bosmia arhinia (Jansz *et al*, 2017). It will be important to determine if disease mutations also impact on DNA damage signaling from telomeres and to what extent this may affect disease pathology.

## Materials and methods

### Cell culture

HeLa cells harboring 11 kb long telomeres as well as the HeLa cells containing an inducible shTRF2 knockdown cassette cell lines were described previously (Grolimund *et al*, 2013). They were used for all transient transfection experiments and to derive *SMCHD1* knockout clones. Cells were maintained at 37°C with 5% CO_2_ in Dulbecco’s modified Eagle’s medium supplemented with 10% FCS and penicillin/streptomycin. For the experiment in Figure 5, WT and *SMCHD1* knockout cells were seeded at 0.5×10^6^ cells per dish. They were irradiated 14 hours post seeding with 3 grays of total γ-irradiation dosage and harvested after the indicated times for Western blot and indirect immunofluorescence analysis.

### Antibodies

The following primary antibodies were used: TRF2 (#05-521, Millipore, mouse, dilution 1:1,000, used for Western blots (WB)), γH2AX (Millipore, #05-636, mouse, dilution 1:1,000, used for WB and IF), hnRNPA1 (4B10, #sc-32301, Santa Cruz Biotechnology, mouse, dilution 1:3,000, used for WB), 53BP1 (#NB100-304, Novus Biologicals, rabbit, dilution 1:2,000, used for IF), phospho-ATM-Ser1981 (#ab81292, Abcam, rabbit, dilution 1:1,000, used for WB and IF), SMCHD1 (#A302-871A, Bethyl Laboratories, N-terminal, rabbit, dilution 1:2,000, used for WB and ChIP), SMCHD1 (#A302-872A, Bethyl Laboratories, C-terminal, rabbit, dilution 1:2,000, used for WB and ChIP), TPP1 (#H00065057-M02, Abnova, rabbit, dilution 1:1,000, used for WB), MRE11 (#NB100-142, Novus Biologicals, rabbit, dilution 1:2,000, used for WB), MDC1 (#ab11171, Abcam, rabbit, dilution 1:1,000, used for WB), normal rabbit IgG (#sc-2027, rabbit, used for ChIP).

### Plasmids

Plasmids containing shRNAs used in this study were prepared by restriction cloning of annealed oligonucleotides into pSUPERpuro or pSUPERblast plasmid backbones (Oligoengine™). The target sequences of the shRNAs were: MRE11 5’-TGAGAACTCTTGGTTTAAC-3’ cloned into pSUPERblast plasmid (Porro *et al*. 2014); TRF2 5’-GCGCATGACAATAAGCAGA-3’ cloned into pSUPERblast and pSUPERpuro plasmid (Porro *et al*. 2014); sh1_SMCHD1 5’-ATTGGATAGCGGGTGATATTA-3’ cloned into pSUPERpuro plasmid; sh2_SMCHD1 5’-TTATTCGAGTGCAACTAATTT-3’ cloned into pSUPERpuro plasmid; shTPP1 5’-GACTTAGATGTTCAGAAAA-3’ cloned into pSUPERblast plasmid (Abreu *et al*. 2010); sh2_MDC1 5’-AGAGGGACAATGATACAAA-3’ cloned into pSUPERblast plasmid; sh3_MDC1 5’-GTCTCCCAGAAGACAGTGA-3’ cloned into pSUPERblast plasmid (Stewart *et al*, 2003) The pSpCas9(BB)-2A-puro plasmid (a generous gift from Dr. Feng Zhang, Addgene plasmid #62988) was used for CRISPR/Cas9 mediated knockout of *SMCHD1*.

### Transfection protocols

For depletion experiments HeLa cells were transfected in 6-well plates at 60-80% confluency using Lipofectamine 2000 according to the manufacturer’s protocol (ThermoFisher, #11668019). Puromycin (conc. 1µg/mL, #ant-pr-1, Invivogen) and blasticidin (conc. 5µg/mL, #ant-bl-1, Invivogen) were added to the media 20-24h after transfection and the cells were expanded on 10cm dishes. Selection with the antibiotics was maintained for 3-5 days. Empty pSuperPURO and pSuperBLAST plasmids were used as control in all the experiments. For the experiment in Figure 4, ATRi (VE-821, Selleckchem, #S8007) was added to the cells 24 hour after addition of the selection antibiotics at 10µM concentration and the cells were maintained with the inhibitor for 4 days.

### Immunoblotting

After harvesting, cells were counted using CASY Cell Counter and Analyzer, cell pellets with equal cell numbers were resuspended in 2x Laemmli buffer (20% glycerol, 4% sodium dodecyl sulphate, 10 mM Tris-Cl pH 6.8, 200 mM Dithiothreitol, 0.05% bromophenol blue) at final concentration of 10 000 cells/µL and boiled for 5min at 95°C. Protein extracts were fractionated on 4-20% Mini-PROTEAN® TGXTM Precast protein gels (Bio-Rad), transferred to a nitrocellulose blotting membrane (AmershamTM ProtranTM, 0.2µm NC, GE Healthcare Life Sciences, #10600001), blocked in 3% BSA/1xPBS/0.1% Tween 20 for 30min and incubated with primary antibody overnight at 4°C. Membranes were then washed 3×5min in 1xPBS/0.1% Tween 20, incubated with anti-mouse or anti-rabbit HRP-conjugated secondary antibody for 30min (anti-mouse IgG HRP conjugate Promega #W402B, anti-rabbit IgG HRP conjugate Promega #W4011, 1:3000) and chemiluminescence was detected using Western Bright ECL spray (Advansta, #K-12049-D50). Detection of TPP1 was performed using a renaturation protocol as described (Loayza & De Lange, 2003).

### Telomere restriction fragment length analysis for detection of single stranded and double stranded telomeric DNA

Genomic DNA was isolated using the Wizard® Genomic DNA Purification kit (Promega, #A1120). Isolated DNA (5 µg) was subjected to digestion with 40U ExoI (New England BioLabs, #M0293S) as control or non-digested and incubated for 8h at 37°C in CutSmart® Buffer in a final volume of 80µL. The samples were then heated at 80°C for 20min to inactivate the ExoI enzyme. Following the inactivation, 20µL of digestion mix containing 125U HinfI (New England BioLabs, #R0155M) and 25U RsaI (New England BioLabs, #R0167L) was added to all the samples (Exo+ and Exo-) and the digestion mix was incubated overnight at 37°C. Digested DNA was loaded on a 1% agarose gel (35µL of the digestion mix was loaded for the Short run and 55µL for the Long run in Figure 2C) and separated by regular gel electrophoresis in 1 × TBE at 3 V cm−1 for 1 h (Short run) and at 1.5 V cm^−1^ for 16h (Long run). Gels were dried for 3h at 50°C, prehybridized at 50°C in Church buffer (1%BSA, 1mM EDTA, 0.5M phosphate buffer pH 7.5, 7%SDS) and hybridized at 50°C overnight to a [32P]-labeled (CCCTAA)n probe (Grolimund *et al*, 2013) for detection of single stranded (ss) telomeric DNA. After hybridization, the gel was rinsed in 4 × SSC and followed by successive 1 h washes at 50°C in 4 × SSC, 4 × SSC/0.5% SDS and 2 × SSC/0.5% SDS and exposed to a sensitive phosphoimager screen overnight. After the image was acquired the gel was denatured with 0.8 M NaOH and 150 mM NaCl, neutralized with 1.5 M NaCl, 0.5 M Tris-HCl pH 7.0, prehybridized in Church buffer at 50° for 1h and incubated with the same probe overnight at 50°C. The gel was again washed and exposed as above and the image was acquired using Amersham™ Typhoon™ Biomolecular imager (GE Healthcare). The images were quantified using Aida Image Analysis software. The single stranded-DNA signal was divided by the total denatured DNA signal in each lane and further normalized to – Dox samples.

### CRISPR/Cas9 gene editing

The CRISPR/Cas9 gene editing system was used to create *SMCHD1* knockout cell lines. To target the *SMCHD1* gene locus (NC_000018.10; gene ID 23347), a region of 200 bp encompassing the ATG in Exon 1 was submitted to the Optimal CRISPR design tool (http://crispr.mit.edu). Three gRNAs with scores higher than 93 were chosen for further experiments (gRNA 1: 5’-CTTGTTTGATCGGCGCGAAA-3’, gRNA2: 5’-GGGGAGCGCTCGGACTACGC-3’, gRNA 3: 5’-GCCGTCCGCCGCTGCCATAT-3’). Complementary oligonucleotides harbouring the guide RNA sequence and BpiI compatible overhangs were synthetized by Microsynth AG. The oligonucleotides were annealed and ligated into a BpiI (Thermo Fisher Scientific, ER1011) digested and dephosphorylated pSpCas9(BB)-2A-puro vector (Addgene, 62988). The resulting constructs were transfected into HeLa cells using LipofectamineTM 2000 (Thermo Fisher Scientific, #11668019). Transfected cells were selected with 1µg/mL of puromycin for 4 days. Single-cell clones were obtained by limiting dilution and were screened for the absence of SMCHD1 by Western blotting using the N-terminal anti-SMCHD1 antibody. To verify the gene editing in positive clones, the PCR products obtained with 2 primers (AV48_SMCHD1_gPCR_F: 5’-AGGAGCGCGTTTGAATCGG-3’, AV47_SMCHD1_gPCR_R 5’-CTTCGCGTACCTGACACACAC-3’) were TOPO-cloned (Thermo Fisher, #450071) and sent for sequencing.

### Telomeric PNA-FISH on metaphase spreads

On the day of harvesting, cells were treated with 0.1 μg/mL demecolcine (Sigma Aldrich Chemie GmbH #D7385-10MG) for 2 h, cells were collected, resuspended in hypotonic solution (0.056 M KCl) and swollen for 7 min. Swollen cells were fixed in methanol:acetic acid (3:1) and stored overnight at 4°C. The next day cell suspensions were dropped onto slides to prepare metaphase spreads, incubated 1min at 70°C in a wet chamber and dried for 16–24 h before FISH. FISH staining of human telomeric DNA was performed as described (Vancevska *et al*, 2017). Slides were rehydrated in 1× PBS for 5 min, treated with 4% formaldehyde in PBS for 5 min, washed 3x with 1xPBS and dehydrated with increasing amounts of ethanol (70%, 95%, 100%). Dehydrated slides were then placed on coverslips containing 70 µL hybridization mix (10 mM Tris-HCl, 2% blocking reagent (Roche, #11096176001), 70% formamide and 0.1 µM Cy3 labeled (CCCTAA)3 PNA probe (PNA Bio, #F1002)) and denatured at 80°C for 3 min in a hybridization oven. Subsequently, the hybridization was allowed to proceed for 3h in a light protected humified chamber at 25 °C. The coverslip was then removed from the slide, washed twice for 15 min in buffer containing 70% formamide and 10 mM Tris-HCl pH 7.4 and 3 times for 15 min with 0.1 M Tris-HCl pH 7.2, 0.15 M NaCl, 0.08% Tween-20. For DNA staining, DAPI was added to 1 µg/ml in the second wash. After the washes slides were stored at 4°C in a dark place until imaging.

### Indirect immunofluorescence and telomeric FISH (IF-FISH)

Indirect immunofluorescence detection of human ATM pS981, 53BP1 and γH2AX followed by telomeric FISH staining was performed as described (Vancevska *et al*. 2017). For detection of ATM pS1981 before crosslinking, cells were fractionated with an ice-cold preextraction buffer containing 0.5% Triton X-100, 20mM HEPES-OH pH 7.5, 50mM NaCl, 3mM MgCl2 and 300mM sucrose for 7min. Subsequently cells were washed with 1xPBS and the same protocol was applied as for the other stainings.

### Chromatin Immuno Precipitation (ChIP)

ChIP protocol for SMCHD1 and γH2AX was performed as described previously (Grolimund *et al*. 2013). Briefly, 10 million cells per condition were harvested and washed with 1xPBS pH 7.4. The cell pellet was then crosslinked in 1mL 1% formaldehyde in 1xPBS pH 7.4 for 15min at RT. Glycine pH 2.5 was added to 125mM to quench the reaction, incubated for 5min and cells were then washed 3x with 1xPBS pH 7.4. Cells were subsequently incubated 5min in 1mL lysis buffer (1% SDS, 10 mM EDTA pH 8.0, 50 mM Tris-Cl pH 8.0, EDTA-free protease inhibitor complex (Roche, #11836170001)), centrifuged 5min at 2,000g and the chromatin enriched pellet was again resuspended in 500µL lysis buffer and subjected to sonication for 30min (30s ON, 30s OFF, total sonication time 15min) using Bioruptor® Twin Diogenode sonicatior (#UCD-400). The sonicated lysate was centrifugated at 20,000g for 15min at 4°C. Per IP 100µL of the cleared lysate was diluted with 9 volumes of IP buffer (1.2 mM EDTA pH 8.0, 1.1% Triton X-100, 16.7 mM Tris pH 8.0, 300 mM NaCl) and incubated with 5µg of the corresponding antibody (normal rabbit IgG, SMCHD1 or γH2AX) and 30µL of preblocked Protein G Sepharose 4 Fast Flow 50% bead slurry (GE Healthcare, #17-0618-01) overnight at 4°C. The beads were then washed with once with wash buffer 1 (0.1% SDS, 1% Triton X-100, 2 mM EDTA pH 8.0, 20 mM Tris pH 8.0, 300 mM NaCl), wash buffer 2 (0.1% SDS, 1% Triton X-100, 2 mM EDTA pH 8.0, 20 mM Tris pH 8.0, 500 mM NaCl), wash buffer 3 (500 mM LiCl, 1% NP-40, 1% Na-deoxycholate, 1 mM EDTA, 10 mM Tris pH 8.0) and twice with wash buffer 4 (1 mM EDTA, 10 mM Tris pH 8.0) at room temperature for 5 min. Elution and crosslink-reversal were performed at 65°C overnight in cross-link reversal buffer (1% SDS, 0.1 M NaHCO3, 0.5 mM EDTA pH 8.0, 20 mM Tris-Cl pH 8.0, 10 μg DNase-free RNase (Roche #11119915001)). For DNA extraction, the QIAquick PCR Purification kit (Qiagen, #28106) was used. Telomeric and Alu-repeat DNA were detected successively using the conditions described before. After the exposure the image was acquired using FujiFilm Fluorescent Image Analyzer FLA-3000 and the image quantification was done using AIDA Image Analyzer software v 4.06.

### Cell cycle analysis

For analysis of the cell cycle distribution of *SMCHD1* KO cells, propidium iodide stained cells were analysed by Fluorescence Activated Cell Sorting (FACS). For each sample 2×10^6^ cells were pelleted, washed in 1xPBS and fixed by dropwise addition of 1mL of ice-cold 70% ethanol and incubated at least 24h hours. Following the fixation, cells were resuspended in 250µL 1xPBS containing 0.2µg/mL of RNAseA and incubated for 15min at 37°C. Cells were then stained by addition of 250µL of 1xPBS containing 80µg/mL propidioum iodide and incubated at 4°C for 10min. Subsequently, cells were passed through a strainer and analysed by FACS on Accuri C6 (BD Biosciences). The percentage of cells in each phase of the cell cycle was determined using the Watson Pragmatic computational model in FlowJo software (TreeStar).

### RT-qPCR for measuring TERRA transcript levels

TERRA was measured as previously described (Feretzaki & Lingner, 2017) with slight modifications. Total RNA was extracted with NucleoSpin® RNA isolation kit (Macherey-Nagel, *#*740955) from 2×10^6^ cells following the manufacturers protocol with 3 DNase treatment steps. cDNA from three biological replicates was synthetized using Invitrogen’s SuperScript III Reverse Transcriptase (#18080044) from 2µg of total RNA. Reaction mixes in total volume of 20µL were prepared as follows: 2µg of total RNA, 0.5mM dNTP mix, 150ng Random primers (Thermo Fisher #48190011), 250ng oligo (dT)15 primer (Promega, #C1101), 1× First-Strand Buffer, 5 mM DTT, 20 U SUPERase IN (Ambion #AM2696) and 200 U SuperScript III RT (200 U/µl) or H2O for no RT-control. The cDNA was then diluted to 40µL and stored at −20°C. Quantitative PCR (qPCR) was performed on an Applied Biosystems 7900HT Fast Real-Time System using Power SYBR Green PCR Master Mix (Applied Biosystems #4368708) in a 384-well reaction plate (Applied Biosystems MicroAmp Optical 384-well reaction Plate with Barcode #4309849). Each sample was prepared in three biological and two technical replicates. The master-mix for each reaction is prepared as follows: 2 µl diluted cDNA, 5 pmol of forward primer, 5 pmol reverse primer, 1× Power SYBR Green PCR Master Mix and H2O to a total volume of 10 µl. qPCR data were analyzed using the relative ΔCt quantification method and GAPDH was used for normalization. Primers used for qPCR are listed in Table 1.

**Table 1.**
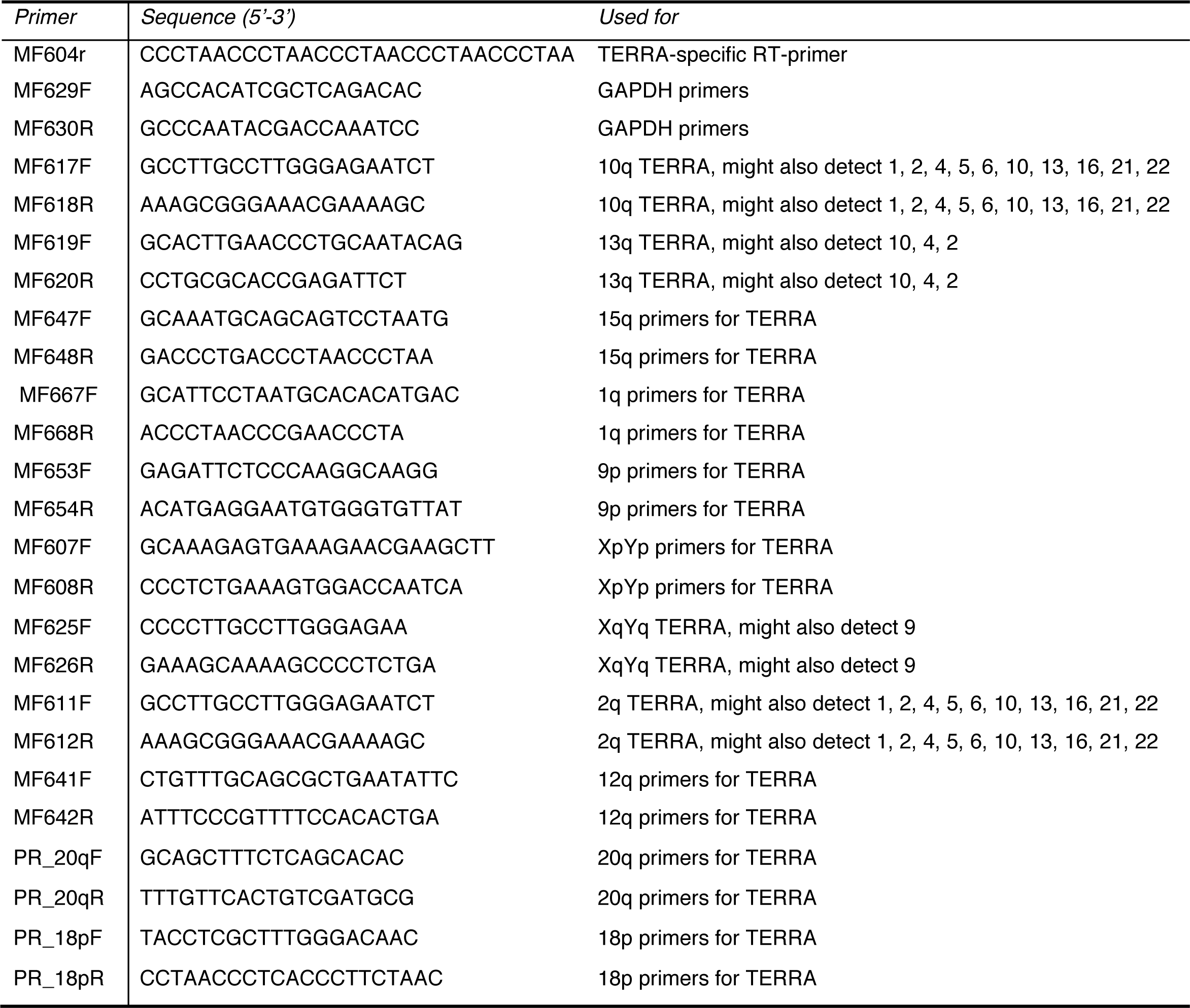
List of oligonucleotides used for RT-qPCR analysis of TERRA expression

## Acknowledgements

We thank Larissa Grolimund for initiating the work on SMCHD1, and Jana Majerska and Galina Glousker for help in establishing the protocol to measure 3’ overhangs. Research in J.L.’s laboratory was supported by the Swiss National Science Foundation (SNSF), the SNSF funded NCCR RNA and disease network, the Swiss Cancer League and EPFL.

## Author Contributions

J.L., A.V. and V.P. designed research, A.V., V.P., M.F. and W.A. carried out experiments, and J.L. and A.V. wrote the paper.

---

**Figure EV1:**
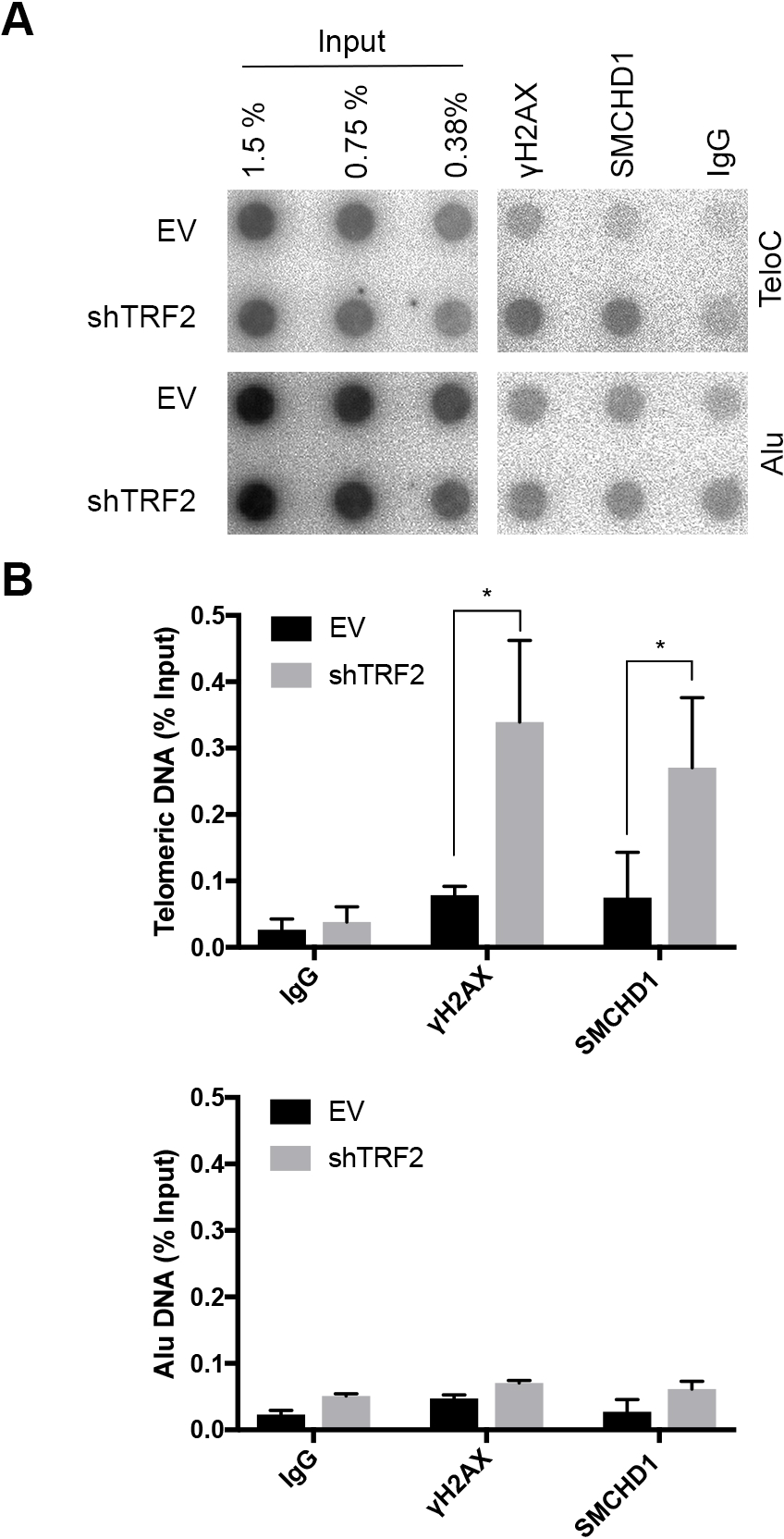
SMCHD1 association with telomeres is increased upon TRF2 removal. A. Telomeric DNA ChIP with antibodies against yH2AX, SMCHD1 and rabbit lgG. Representative dot blot images of precipitated DNA detected with a (CCCTAA)n or Alu probe. ChlP s were performed in Hela cells transfected with shTRF2 or Empty Vector (EV) control.
B. Bar graph for quantification of yH2AX, SMCHD1 and rabbit lgG binding to telomeric or Alu DNA as assesed by ChlP. The bars represent average values from three independent experiments for telomeric DNA, and two independent experiments for Alu DNA. Error bars represent the standard deviation. P-values were calculated by unpaired two-tailed Student’s t-test (*) P< 0.05.

**Figure EV2:**
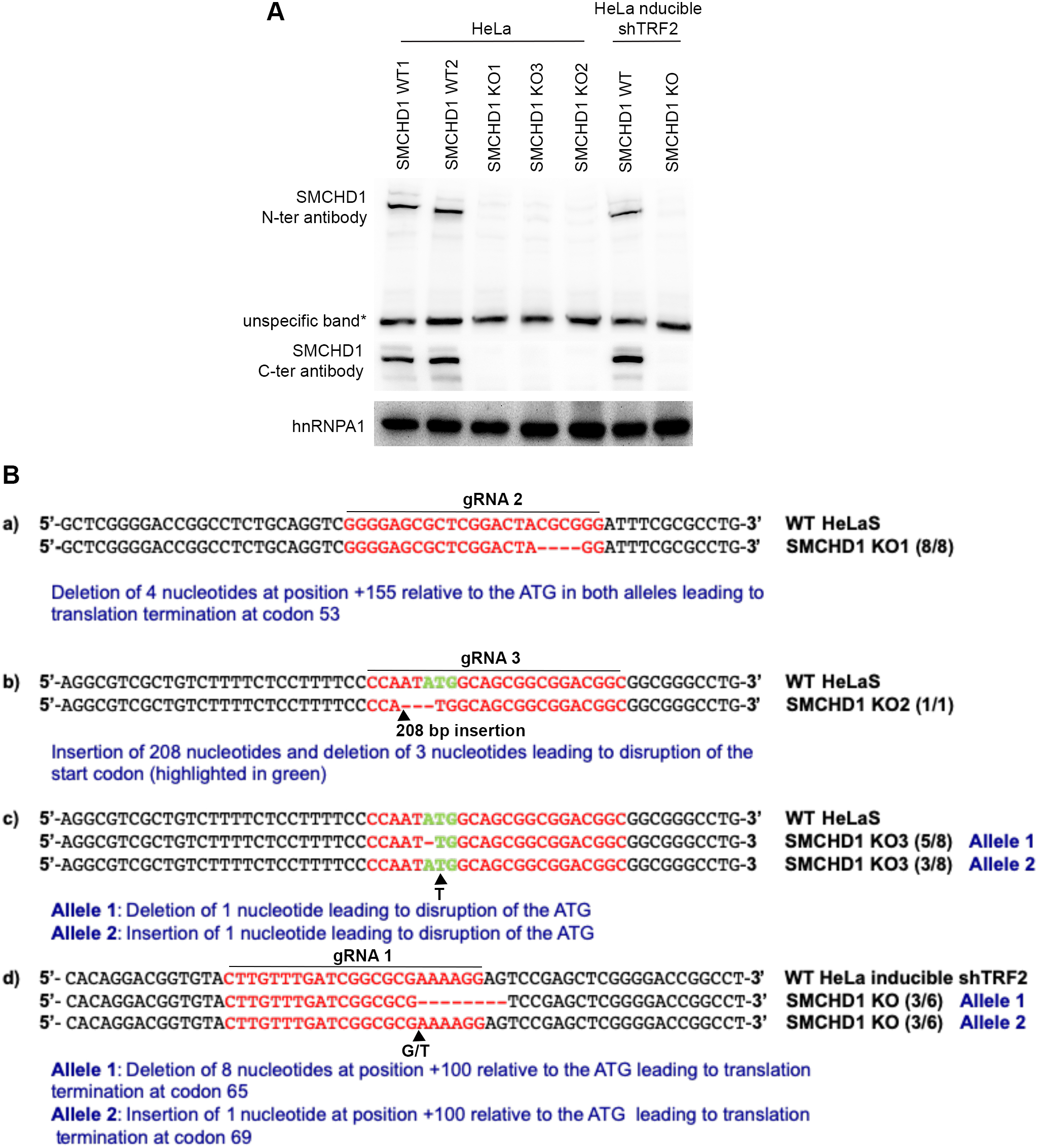
Generation of SMCHD1 knockout cells using CR1SPR-Cas9 technology. A. Western Blot detection of SMCHD1 with an antibody raised against the N-terminus (aa213-aa300) and the C-terminus of the protein (aa1955-2005) in *SMCHD1* knockout single cell clones of Hela and Hela inducible shTRF2 cell lines.
B. Sequence analysis of edited alleles in *SMCHD1* knockout single cell clones.

**Figure EV3:**
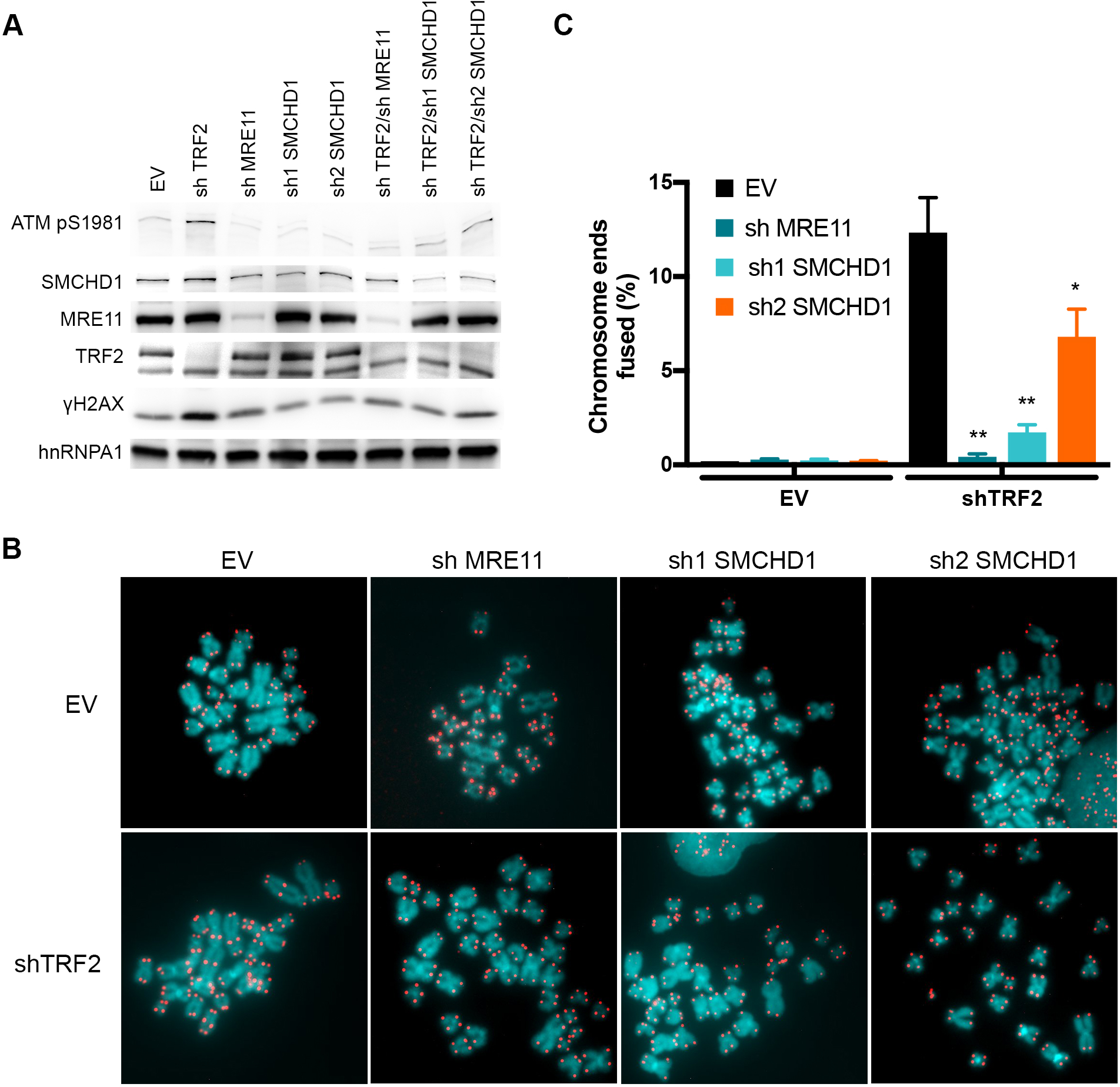
SMCHD1 stimulates c-NHEJ at TRF2 depleted telomeres. A. Western Blot detection of ATM pS1981, SMCHD1, MRE11, TRF2, and yH2AX, in Hela cells transfected with the indicated shRNA plasmids (shTRF2, shMRE11, sh1SMCHD1, sh2SMCHD1, shTRF2/shMRE11, shTRF2/sh1 SMCHD1, shTRF2/sh2 SMCHD1) and empty vector (EV) control.
B. Metaphase spreads from Hela cells transfected with the indicated shRNA plasmids (shTRF2, shMRE11, sh1SMCHD1, sh2SMCHD1, shTRF2/shMRE11, shTRF2/sh1 SMCHD1, shTRF2/sh2 SMCHD1) and empty vector (EV) control. Telomeric signals were detected with C y3-(CCC TAA)3 and are false colored in red.
C. Quantification of telomere fusions in Hela cells transfected with the indicated shRNAs and EV control. Bars represent average numbers of chromosome ends fused in 3 independent experiments with SDs (>6500 telomeres counted/ condition/ experiment). (**) P < 0.01; (*) P < 0.05, unpaired two-tailed

**Figure EV4:**
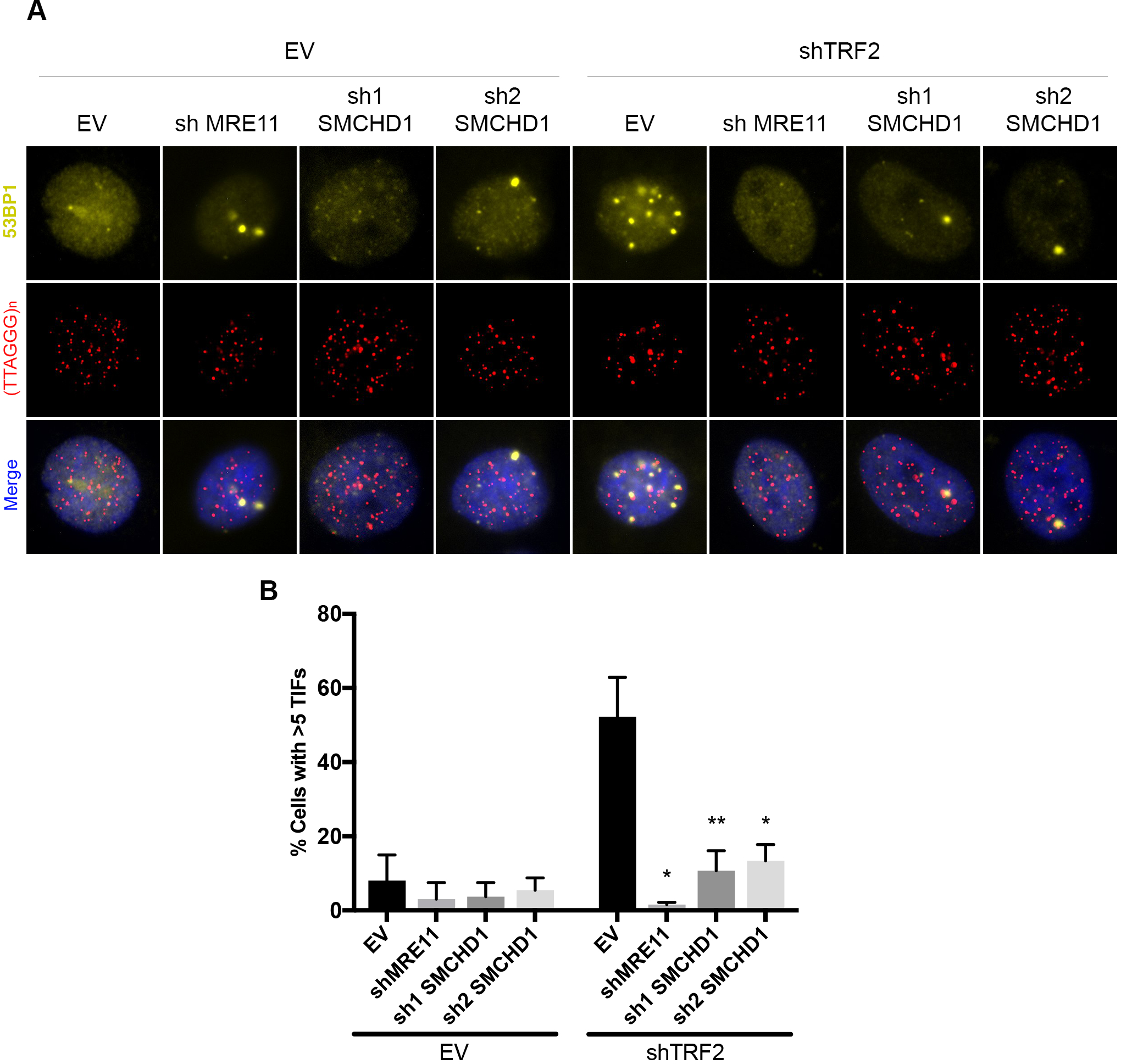
SMCHD1 promotes DNA damage signaling at TRF2 depleted telomeres. A. Representative images for detection of 53BP1 at telomeres in Hela cells transfected with the indicated shRNA plasmids. lmmunofluoresence (IF) for 53BP1 (yellow) was combined with telomeric (CCCTAA)_3_-FISH (red) and the DNA was stained with DAPI.
B. Quantification of the number of cells containing >5 Telomere dysfunction Induced Foci (T IFs) detected as in (B). Data represent mean of 3 independent experiments± SD (>100 cells/condition/experiment). (*) P<0.05, (**) P< 0.01, unpaired two tailed Student’s t-test.

**Figure EV5:**
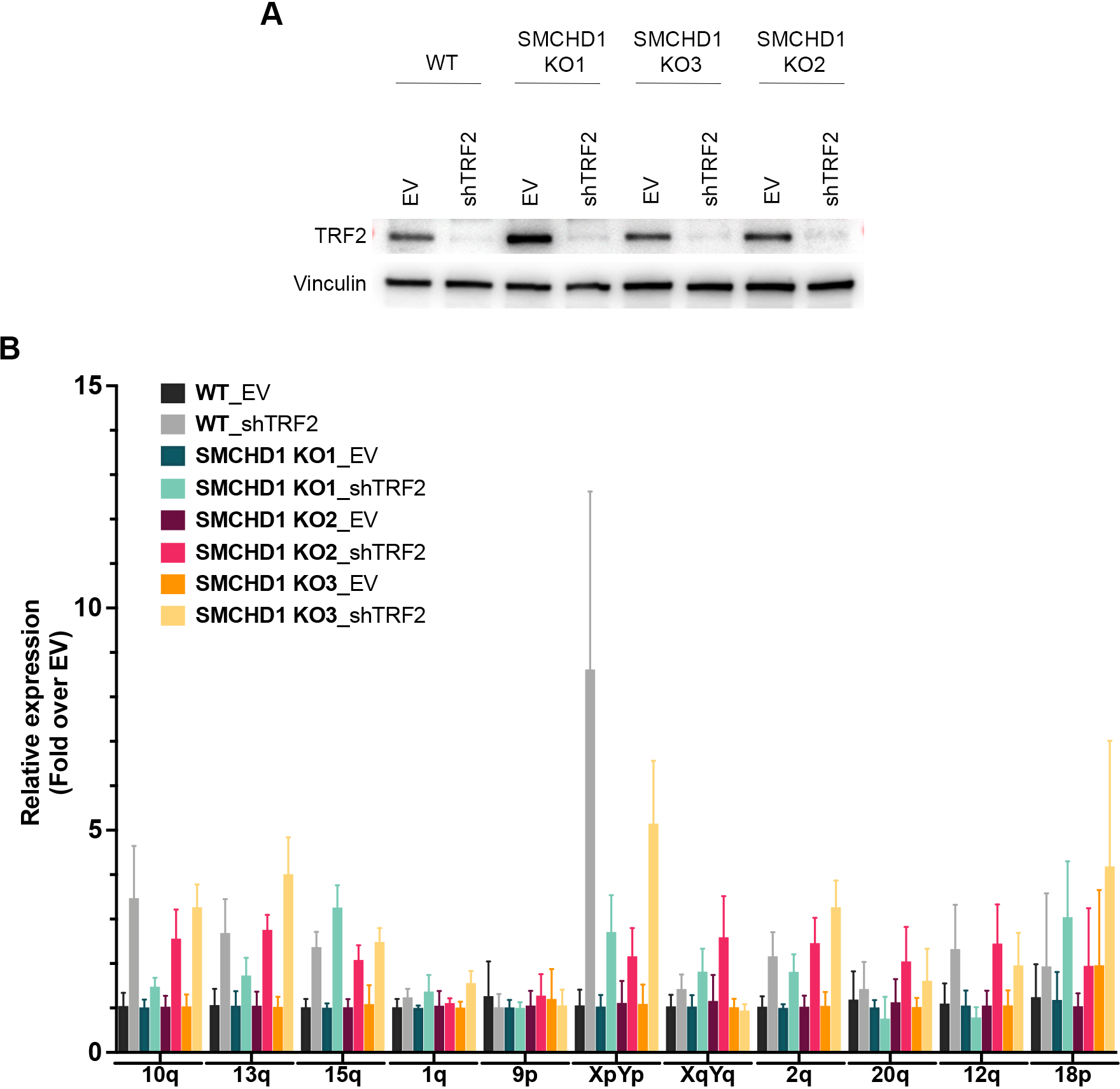
Effects of SMCHD1 KO in shTRF2 depleted Hela cells on TERRA expression A. Western Blot detection of SMCHD1, TRF2 and Vinculin in WT and SMCHD1 KO Hela cells transfected with shTRF2 plasmid or EV control.
B. TERRA quantification by RT-qPCR using primers specfic for the indicated subtelomeric sequences (10q, 13q, 15q, 1q, 9p, XpYp, XqYq, 2q, 20q, 12q, 18p). Bars represent average expression levels normalized to EV in each indicated cell line. SDs were obtained from three biological and two techical replicates.

**Figure EV6:**
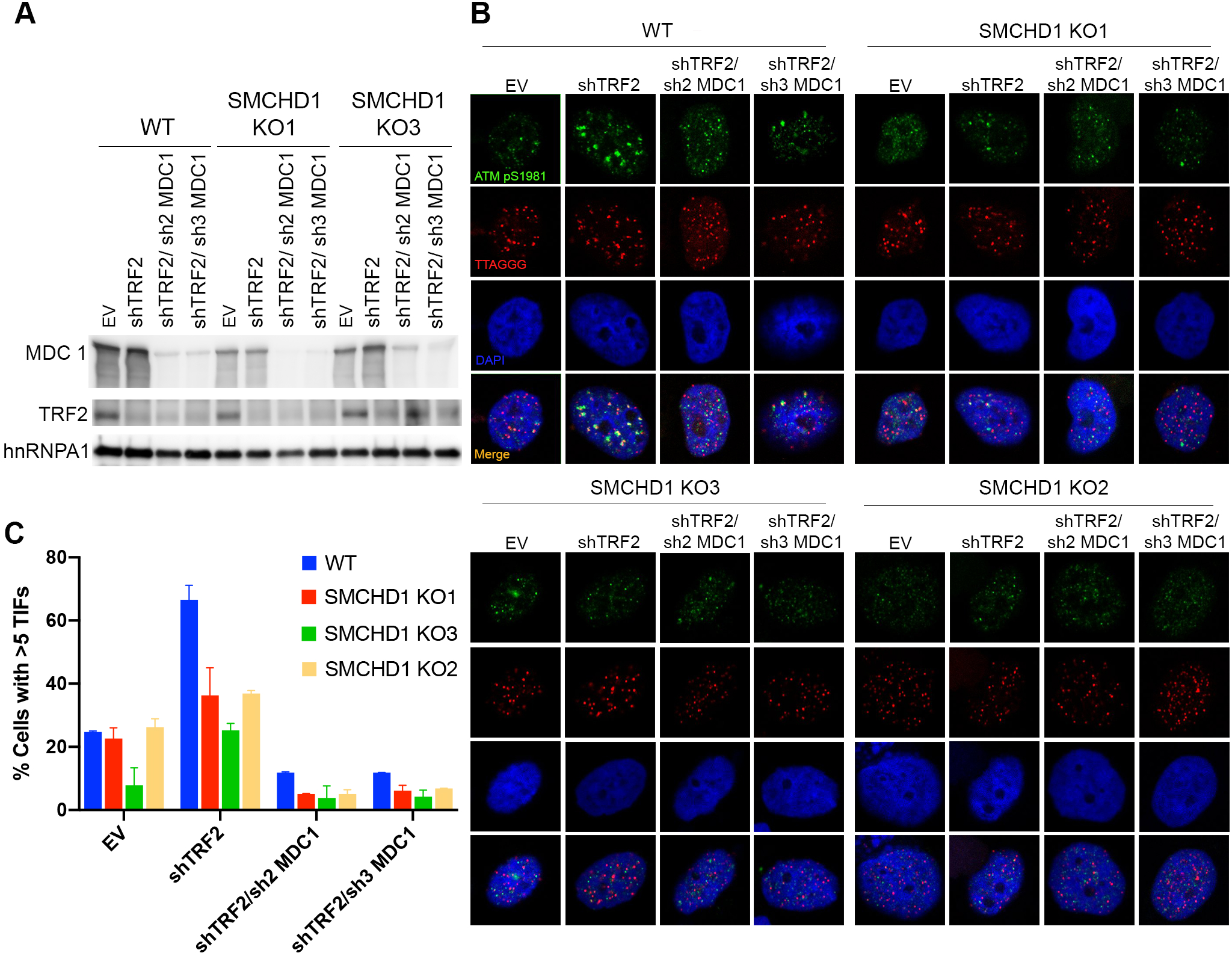
Analysis of epistatic relationship between MDC1 and SMCHD1. A. Western Blot detection of MDC1, TRF2 and hnRNPA1 in WT or SMCHD1 KO Hela cells transfected with the indicated shRNA plasmids (shTRF2, shTRF2/sh2 MDC1, shTRF2/sh3 MDC1) and EV control.
B. Representative images for detection of ATM pS9181 at telomeres in Wild Type (WT) and SMCHD1 KO Hela cells transfected with idicated shRNA plasmids and empty vector (EV) control. lmmunofluoresence (IF) for ATM pS1981 (green) was combined with telomeric (CCCTAA)3-F1SH (red) and the DNA was stained with DAPI.
C. Quantification of the number of cells containing >5 Telomere dysfunction Induced Foci (TIFs) detected as in B. Data represent mean of 2 independent experiments± SD (>50 cells/condition/experiment).

## References

Bartocci C, Diedrich JK, Ouzounov I, Li J, Piunti A, Pasini D, Yates JR & Lazzerini Denchi E (2014) Isolation of chromatin from dysfunctional telomeres reveals an important role for Ring1b in NHEJ-mediated chromosome fusions. Cell Rep 7: 1320–1332

Blewitt ME, Gendrel A-V, Pang Z, Sparrow DB, Whitelaw N, Craig JM, Apedaile A, Hilton DJ, Dunwoodie SL, Brockdorff N, Kay GF & Whitelaw E (2008) SmcHD1, containing a structural-maintenance-of-chromosomes hinge domain, has a critical role in X inactivation. Nat. Genet. 40: 663–669

Brideau NJ, Coker H, Gendrel A-V, Siebert CA, Bezstarosti K, Demmers J, Poot RA, Nesterova TB & Brockdorff N (2015) Independent Mechanisms Target SMCHD1 to Trimethylated Histone H3 Lysine 9-Modified Chromatin and the Inactive X Chromosome. Mol. Cell. Biol. 35: 4053–4068

Celli GB & de Lange T (2005) DNA processing is not required for ATM-mediated telomere damage response after TRF2 deletion. Nat. Cell Biol. 7: 712–718

Cesare AJ, Hayashi MT, Crabbe L & Karlseder J (2013) The telomere deprotection response is functionally distinct from the genomic DNA damage response. Mol. Cell 51: 141–155

Ciccia A & Elledge SJ (2010) The DNA damage response: making it safe to play with knives. Mol. Cell 40: 179–204

Coker H & Brockdorff N (2014) SMCHD1 accumulates at DNA damage sites and facilitates the repair of DNA double-strand breaks. J. Cell. Sci. 127: 1869–1874

Déjardin J & Kingston RE (2009) Purification of proteins associated with specific genomic Loci. Cell 136: 175–186

Denchi EL & de Lange T (2007) Protection of telomeres through independent control of ATM and ATR by TRF2 and POT1. Nature 448: 1068–1071

Deng Y, Guo X, Ferguson DO & Chang S (2009) Multiple roles for MRE11 at uncapped telomeres. Nature 460: 914–918

Dimitrova N & de Lange T (2006) MDC1 accelerates nonhomologous end-joining of dysfunctional telomeres. Genes Dev. 20: 3238–3243

Doksani Y, Wu JY, de Lange T & Zhuang X (2013) Super-resolution fluorescence imaging of telomeres reveals TRF2-dependent T-loop formation. Cell 155: 345–356

d’Adda di Fagagna F, Reaper PM, Clay-Farrace L, Fiegler H, Carr P, Von Zglinicki T, Saretzki G, Carter NP & Jackson SP (2003) A DNA damage checkpoint response in telomere-initiated senescence. Nature 426: 194–198

Feretzaki M & Lingner J (2017) A practical qPCR approach to detect TERRA, the elusive telomeric repeat-containing RNA. Methods 114: 39–45

Frescas D & de Lange T (2014) Binding of TPP1 protein to TIN2 protein is required for POT1a,b protein-mediated telomere protection. J. Biol. Chem. 289: 24180–24187

Gdula MR, Nesterova TB, Pintacuda G, Godwin J, Zhan Y, Ozadam H, McClellan M, Moralli D, Krueger F, Green CM, Reik W, Kriaucionis S, Heard E, Dekker J & Brockdorff N (2019) The non-canonical SMC protein SmcHD1 antagonises TAD formation and compartmentalisation on the inactive X chromosome. Nat Commun 10: 30

Goodarzi AA, Noon AT, Deckbar D, Ziv Y, Shiloh Y, Löbrich M & Jeggo PA (2008) ATM signaling facilitates repair of DNA double-strand breaks associated with heterochromatin. Mol. Cell 31: 167–177

Grolimund L, Aeby E, Hamelin R, Armand F, Chiappe D, Moniatte M & Lingner J (2013) A quantitative telomeric chromatin isolation protocol identifies different telomeric states. Nat Commun 4: 2848

Jansz N, Chen K, Murphy JM & Blewitt ME (2017) The Epigenetic Regulator SMCHD1 in Development and Disease. Trends Genet. 33: 233–243

Jansz N, Keniry A, Trussart M, Bildsoe H, Beck T, Tonks ID, Mould AW, Hickey P, Breslin K, Iminitoff M, Ritchie ME, McGlinn E, Kay GF, Murphy JM & Blewitt ME (2018) Smchd1 regulates long-range chromatin interactions on the inactive X chromosome and at Hox clusters. Nat. Struct. Mol. Biol. 25: 766–777

Jones RE, Oh S, Grimstead JW, Zimbric J, Roger L, Heppel NH, Ashelford KE, Liddiard K, Hendrickson EA & Baird DM (2014) Escape from telomere-driven crisis is DNA ligase III dependent. Cell Rep 8: 1063–1076

Karlseder J, Broccoli D, Dai Y, Hardy S & de Lange T (1999) p53- and ATM-dependent apoptosis induced by telomeres lacking TRF2. Science 283: 1321–1325

Karlseder J, Smogorzewska A & de Lange T (2002) Senescence induced by altered telomere state, not telomere loss. Science 295: 2446–2449

de Lange T (2018) Shelterin-Mediated Telomere Protection. Annu. Rev. Genet. 52: 223–247

Lazzerini-Denchi E & Sfeir A (2016) Stop pulling my strings – what telomeres taught us about the DNA damage response. Nat. Rev. Mol. Cell Biol. 17: 364–378

Loayza D & De Lange T (2003) POT1 as a terminal transducer of TRF1 telomere length control. Nature 423: 1013–1018

Maciejowski J & de Lange T (2017) Telomeres in cancer: tumour suppression and genome instability. Nat. Rev. Mol. Cell Biol. 18: 175–186

Matsuoka S, Ballif BA, Smogorzewska A, McDonald ER, Hurov KE, Luo J, Bakalarski CE, Zhao Z, Solimini N, Lerenthal Y, Shiloh Y, Gygi SP & Elledge SJ (2007) ATM and ATR substrate analysis reveals extensive protein networks responsive to DNA damage. Science 316: 1160–1166

McClintock B (1941) The Stability of Broken Ends of Chromosomes in Zea Mays. Genetics 26: 234–282

Muller HJ (1938) The remaking of chromosomes. Collecting Net. 13: 181–198

Nozawa R-S, Nagao K, Igami K-T, Shibata S, Shirai N, Nozaki N, Sado T, Kimura H & Obuse C (2013) Human inactive X chromosome is compacted through a PRC2-independent SMCHD1-HBiX1 pathway. Nat. Struct. Mol. Biol. 20: 566–573

Okamoto K, Bartocci C, Ouzounov I, Diedrich JK, Yates JR & Denchi EL (2013) A two-step mechanism for TRF2-mediated chromosome-end protection. Nature 494: 502–505

Panier S & Durocher D (2013) Push back to respond better: regulatory inhibition of the DNA double-strand break response. Nat. Rev. Mol. Cell Biol. 14: 661–672

Paull TT (2015) Mechanisms of ATM Activation. Annu. Rev. Biochem. 84: 711–738

Porro A, Feuerhahn S, Delafontaine J, Riethman H, Rougemont J & Lingner J (2014a) Functional characterization of the TERRA transcriptome at damaged telomeres. Nat Commun 5: 5379

Porro A, Feuerhahn S & Lingner J (2014b) TERRA-Reinforced Association of LSD1 with MRE11 Promotes Processing of Uncapped Telomeres. Cell Reports 6: 765–776

Rai R, Hu C, Broton C, Chen Y, Lei M & Chang S (2017) NBS1 Phosphorylation Status Dictates Repair Choice of Dysfunctional Telomeres. Mol. Cell 65: 801–817.e4

Reaper PM, Griffiths MR, Long JM, Charrier J-D, Maccormick S, Charlton PA, Golec JMC & Pollard JR (2011) Selective killing of ATM- or p53-deficient cancer cells through inhibition of ATR. Nat. Chem. Biol. 7: 428–430

van Ruiten MS & Rowland BD (2018) SMC Complexes: Universal DNA Looping Machines with Distinct Regulators. Trends Genet. 34: 477–487

Shay JW & Wright WE (2011) Role of telomeres and telomerase in cancer. Semin. Cancer Biol. 21: 349–353

Smogorzewska A, van Steensel B, Bianchi A, Oelmann S, Schaefer MR, Schnapp G & de Lange T (2000) Control of human telomere length by TRF1 and TRF2. Mol. Cell. Biol. 20: 1659–1668

Stewart GS, Wang B, Bignell CR, Taylor AMR & Elledge SJ (2003) MDC1 is a mediator of the mammalian DNA damage checkpoint. Nature 421: 961–966

Sun Y, Jiang X, Xu Y, Ayrapetov MK, Moreau LA, Whetstine JR & Price BD (2009) Histone H3 methylation links DNA damage detection to activation of the tumour suppressor Tip60. Nat. Cell Biol. 11: 1376–1382

Syed A & Tainer JA (2018) The MRE11-RAD50-NBS1 Complex Conducts the Orchestration of Damage Signaling and Outcomes to Stress in DNA Replication and Repair. Annu. Rev. Biochem. 87: 263–294

Takai H, Smogorzewska A & de Lange T (2003) DNA damage foci at dysfunctional telomeres. Curr. Biol. 13: 1549–1556

Tang M, Li Y, Zhang X, Deng T, Zhou Z, Ma W & Songyang Z (2014) Structural maintenance of chromosomes flexible hinge domain containing 1 (SMCHD1) promotes non-homologous end joining and inhibits homologous recombination repair upon DNA damage. J. Biol. Chem. 289: 34024–34032

Van Ly D, Low RRJ, Frölich S, Bartolec TK, Kafer GR, Pickett HA, Gaus K & Cesare AJ (2018) Telomere Loop Dynamics in Chromosome End Protection. Mol. Cell 71: 510–525.e6

Vancevska A, Douglass KM, Pfeiffer V, Manley S & Lingner J (2017) The telomeric DNA damage response occurs in the absence of chromatin decompaction. Genes Dev. 31: 567–577

Wang C-Y, Jégu T, Chu H-P, Oh HJ & Lee JT (2018) SMCHD1 Merges Chromosome Compartments and Assists Formation of Super-Structures on the Inactive X. Cell 174: 406–421.e25

Zhu XD, Küster B, Mann M, Petrini JH & de Lange T (2000) Cell-cycle-regulated association of RAD50/MRE11/NBS1 with TRF2 and human telomeres. Nat. Genet. 25: 347–352

Zou L & Elledge SJ (2003) Sensing DNA damage through ATRIP recognition of RPA-ssDNA complexes. Science 300: 1542–1548

